# EveD, a vesicle-associated rhizobial protein shaping nodulation competitiveness and host genotype responses

**DOI:** 10.64898/2026.11.12.766089

**Authors:** Diego García-Rodríguez, Paula Ayala-García, Francisco Pérez-Montaño, Sebastián Acosta-Jurado, José María Vinardell, José Manuel Borrero-de Acuña, Francisco Javier Ollero, Francisco Javier López-Baena, Irene Jiménez-Guerrero

## Abstract

Bacterial extracellular vesicles (bEVs) are an ancestral secretion route that delivers cargoes to host cells. Although it is known that rhizobial bEVs transport Nod factors and other bacterial components, their role in protein delivery during symbiosis remains unexplored. Here we characterise EveD, a large soluble repeat-containing protein of *Sinorhizobium fredii* HH103. Its export is independent of the type III secretion system, and EveD is recovered only in bEV-containing fractions. *eveD* expression is not induced by genistein or soybean seed exudates and does not follow the canonical NodD1-TtsI pathway, but partially depends on the symbiotic regulator SyrM. When expressed in plant cells, EveD localises mainly to the cytoplasm, associates with endoplasmic reticulum-like structures and occasionally occurs in the nucleus. The *eveD* mutant shows increased nodule occupancy relative to the parental strain in *Glycine max* cv. Williams. EveD modulates early responses in wild *Glycine soja* accessions in a genotype-dependent manner, constraining pseudonodule organogenesis in JL2 while promoting it in CH2. These findings identify EveD as a bEV-associated rhizobial protein that shapes bacterial competitiveness and genotype-dependent symbiotic responses, and highlight bEVs as an additional route for rhizobial protein export in symbiosis with legumes.

**Highlight:** EveD is associated with rhizobial extracellular vesicles and influences bacterial competitiveness and genotype-dependent pseudonodule organogenesis.

## Introduction

Extracellular vesicles (EVs) are nanoscale lipid-bound structures, typically ranging from 20 to 400 nm in diameter, produced by organisms from all domains of life (Schwechheimer and Kuehn, 2015; Gill *et al*., 2019; Toyofuku *et al*., 2023). Bacterial EVs (bEVs) constitute a sophisticated secretion strategy that expands the classical view of bacterial molecular export. Recently proposed as the Type 0 Secretion System (T0SS; Guerrero-Mandujano *et al*., 2017), bEV-mediated secretion is considered an evolutionarily conserved mechanism that enables bacteria to package and deliver diverse biological cargoes, including proteins, lipids, nucleic acids, and metabolites (Toyofuku *et al*., 2019; 2023).

In Gram-negative bacteria, bEVs are mainly generated through either outer/outer-inner membrane/s blebbing or explosive cell lysis, resulting in vesicles derived from the outer membrane or, in some cases, containing components from both the outer and inner membranes (Turnbull *et al*., 2016; Toyofuku *et al*., 2017; 2019; 2023). By enabling safe transport and targeted delivery of biologically active molecules, bEVs contribute to diverse biological processes, including nutrient acquisition, microbial competition, host colonisation, and immune modulation (Kadurugamuwa and Beveridge, 1996; Fulsundar *et al*., 2014; Toyofuku *et al*., 2017; 2023; Li *et al*., 2022; Wang *et al*., 2023). Importantly, bEVs can concentrate bioactive molecules within its cargo, a process termed quantal secretion, which ensures the delivery of biologically relevant doses while preserving molecular activity over long distances (Toyofuku *et al*., 2017). Together, these properties establish bEVs as versatile delivery platforms that allow bacteria to modulate their environmental adaptation and interact with other organisms.

Bacterial bEV production and cargo composition are dynamic processes modulated by environmental and host-derived signals, suggesting that vesicle formation represents a regulated strategy to adapt bacterial communication to specific ecological conditions (McBroom *et al*., 2006; Bomberger *et al*., 2011; Schwechheimer and Kuehn, 2015).

In the particular case of plant-associated bacteria, most studies to date have focused on pathogenic bacteria, in which bEVs have been shown to contribute to host colonisation and disease development by transporting virulence-associated proteins, toxins, enzymes, and regulatory molecules capable of modulating plant immunity (Bahar *et al*., 2016; Katsir and Bahar, 2017; Rudnicka *et al*., 2022; Wang *et al*., 2023). In contrast, the role of bEVs from beneficial plant-associated bacteria remains comparatively unexplored (Maqbool *et al*., 2026). Among these bacteria, rhizobia, soil-borne α- and β-proteobacteria, represent a particularly relevant system to investigate bEV-mediated communication, as their highly specialised symbiosis with legumes relies on a tightly regulated molecular dialogue between both partners. This complex interaction ultimately leads to the formation of root nodules by the plant, specialised organs where rhizobia are intracellularly hosted, differentiate into nitrogen-fixing bacteroids and provide reduced nitrogen to the plant in exchange for carbon compounds (Oldroyd, 2013; Roy *et al*., 2020). The initial recognition between compatible rhizobia and legume is mediated by the perception of plant-derived flavonoids and the production and secretion of bacterial Nod factors (NFs). Flavonoids released by legume roots activate the bacterial transcriptional regulator NodD, which subsequently binds to conserved promoter sequences called *nod* boxes located upstream of nodulation genes, inducing their expression. NFs are lipochitooligosaccharidic molecules recognised by plant receptors that trigger a signalling cascade that promotes root hair deformation, infection thread formation, and nodule organogenesis (Oldroyd, 2013; Zipfel and Oldroyd, 2017).

Beyond their secretion as soluble signals, NFs can also be transported via bEVs, providing a mechanism to protect these labile signalling molecules from degradation and facilitate their interaction with plant roots (Taboada *et al*., 2019ab; Li *et al*., 2022; Moreno de Castro *et al*., 2026). In *Sinorhizobium fredii* HH103, NF-containing bEVs promote root hair deformation, activate legume nodulation genes and repress plant defence genes, aiding early recognition and immune modulation (Li *et al*., 2022). In fact, genistein-induced bEVs were shown to stimulate nodule primordia formation and enhance soybean symbiotic performance in field trials when supplementing commercial inoculants, which is why these vesicles have been termed Plant Growth-Promoting Vesicles (PGPVs) (Moreno de Castro *et al*., 2026).

bEVs may also contribute to this molecular dialogue by transporting additional bacterial components involved in host recognition. For example, the outer membrane protein MhOpa22 of *Mesorhizobium huakuii* exerts its symbiotic function via bEVs and interacts with the plant germin-like protein AsGLP1 in *Astragalus sinicus* (Zeng *et al*., 2023; Li *et al*., 2025). In later stages of the symbiosis, host and bacterial EVs also traffic within the nodule peribacteroid space, where they regulate conserved processes despite differential EV-associated proteomes, indicating an EV-mediated dialogue across legume species (Ayala-García *et al*., 2025).

In *S. fredii* HH103, expression of symbiotic genes is governed by a complex regulatory network in which NodD1 acts as the primary activator in response to host-derived flavonoids (Pérez-Montaño *et al*., 2016). Additional regulators including NodD2, NolR, and SyrM, further contribute to fine-tune symbiotic gene expression and NFs production (Pérez-Montaño *et al*., 2016; Acosta-Jurado *et al*., 2019; 2020; Navarro-Gómez *et al*., 2023; Jiménez-Guerrero *et al*., 2025). Moreover, the transcriptional activator TtsI links NodD1-dependent flavonoid signalling to the induction of the Type III secretion system (T3SS), a macromolecular apparatus that delivers proteins directly into eukaryotic host cells (Galán and Collmer, 1999). Its regulatory cascade, structural components, and complete repertoire of secreted effectors (T3Es) have been extensively characterised in this strain (Rodrigues *et al*., 2007; López-Baena *et al*., 2008; 2009; Jiménez-Guerrero *et al*., 2015a; 2015b; 2017; 2019; 2020; 2025; Pérez-Montaño *et al*., 2016). Besides, while some *S. fredii* HH103 T3Es, such as NopI, NopL, and NopP, appear to be associated with symbiosis-specific functions, others, including NopD, NopM, and NopT, share homology with effectors from eukaryotic pathogens, suggesting conserved mechanisms of host manipulation (Jiménez-Guerrero *et al*., 2022; Camuel *et al*., 2026).

In this study, we identified and studied the Extracellular vesicle candidate effector D (EveD) protein, a previously uncharacterised large protein from *S. fredii* HH103 with an extended α-helical tandem repeat architecture, initially identified through its sequence similarity to the *Xanthomonas* transcription activator-like (TAL) effector AvrBs3. We demonstrated that EveD is an bEV-associated protein regulated independently of the canonical T3SS pathway, which mainly localises to the host cytoplasm and modulates *S. fredii* HH103-soybean symbiosis in a host-dependent manner. Our findings suggest that EveD may represent a previously unrecognised family of rhizobial type 0 specific effectors.

## Materials and Methods

### Bacterial strains and growth conditions

Bacterial strains used in this work are listed in Supplementary Table S1. *S. fredii* HH103 was grown at 28 °C on tryptone yeast (TY) (Beringer, 1974), yeast extract mannitol (YM) (Vincent, 1970) or minimal (MM) (Robertsen *et al*., 1981) media. *Escherichia coli* and *Agrobacterium-*derived strains were cultured in lysogeny broth (LB) (Lennox LB 5 g l^-1^ NaCl) supplemented with 2% w/v agar, when necessary (Sambrook *et al*., 1989) at 37 °C and 28 °C, respectively. When required, the media were supplemented with the antibiotics gentamicin (Gm, 10 µg ml^-1^ for *E. coli* and *S. fredii* HH103), kanamycin (Km, 30 µg ml^-1^ for *E. coli,* 60 µg ml^-1^ for *S. fredii* HH103, and 50 µg ml^-1^ for *A. tumefaciens*), nalidixic acid (Nx, 20 µg ml^-1^ for *E. coli*), spectinomycin (Spc, 100 µg ml^-1^ for *E. coli* and *A. tumefaciens* K599, and 50 µg ml^-1^ for *S. fredii* HH103), tetracycline (Tc, 10 µg ml^-1^ for *E. coli*, and 5 µg ml^-1^ for *S. fredii* HH103) and rifampicin (50 µg ml^-1^ for *S. fredii* HH103 and *A. tumefaciens* GV3101). Genistein, a *nod* gene-inducing flavonoid, was dissolved in ethanol at a concentration of 1 µg ml^-1^ to obtain a final concentration of 3.7 µM. To obtain soybean exudates, seeds were surface-sterilised and imbibed in sterile water for 48 h at 4 °C (200 mg of seeds ml^-1^). The resulting seed exudates were filtered through 0.45 µm cellulose acetate syringe filters and stored at -20 °C until use, and diluted to a final concentration of 16% (v/v) for subsequent experiments (Pérez-Montaño *et al*. 2013).

### Construction of plasmids and bacterial strains

Plasmids and primers used in this study are listed in Tables S1 and S2, respectively. To construct the *S. fredii* HH103 *eveD* mutant, a 5’fragment of this gene including the promoter region was amplified using primers eveDext_BamHI and eveDint_omega_XbaIR. The PCR product was digested with *Bam*HI and *Xba*I, and cloned into the pK18*mob* suicide vector (Schäfer *et al*., 1994), which was previously digested with the same enzymes, obtaining plasmid pK18*mob*::*eveD*. In parallel, the 2 kb Ω interposon from plasmid pHP45Ω (Prentki and Krisch, 1984) was amplified by PCR using primers InOmega_ClaIF and InOmega2_ClaIR, digested with *ClaI*, and inserted into the *Cla*I site within the *eveD* gene in pK18*mob*::*eveD,* generating the plasmid pK18*mob*::*eveD*::Ω. This construct was employed to generate the *S. fredii* HH103 *eveD*::Ω mutant strain via double homologous recombination. Homogenisation was confirmed by Southern blot, PCR and sequencing. For Southern blot hybridisation, DNA was blotted onto Hybond-N nylon membranes (Amersham, UK), and the DIG-DNA labeling method from Roche (Switzerland) was used following the manufacturer’s instructions. The correct insertion of the Ω interposon into the *eveD* gene was assessed by Southern blot analysis using a specific probe (Supplementary Fig. S1A). The absence of EveD in the *S. fredii* HH103 *eveD*::Ω mutant was evaluated by Western blot analysis using a specific anti-EveD antibody (Supplementary Fig. S1B). Polyclonal antibodies were custom-generated by Davids Biotechnologie GmbH (Germany) using an *in silico*-predicted, highly immunogenic EveD peptide synthesised *de novo* (1 rabbit; 4 protein injections - 500, 125, 125 and 125 μg – and 2 bleedings).

Complementation of the *eveD* mutation could not be achieved by direct amplification of the full-length gene due to the inability to obtain a large PCR product. Therefore, an alternative strategy was employed based on amplification of the gene in two overlapping fragments, followed by assembly through restriction digestion and ligation. Both fragments were amplified by PCR from genomic DNA of the *S. fredii* HH103 parental strain. The first fragment (∼3.5 kb) was obtained using the primers eveDext_BamHI and eveD_IntComp_R, introducing a *BamH*I restriction site. The second fragment (∼2.2 kb) was amplified using the primers eveD_IntComp_F and eveD_XbaI_HA_R, introducing a C-terminal haemagglutinin (HA) epitope coding sequence and an *Xba*I restriction site. The HA tag was used to verify EveD synthesis by immunodetection. The two fragments share an overlapping region containing the native *Nco*I restriction site. Then, the first fragment was digested with *Nco*I and *Bam*HI, while the second fragment was digested with *Nco*I and *Xba*I. Subsequently, both fragments were ligated simultaneously into plasmid pBBR1-MCS5 (Kovach *et al*., 1995), previously linearised with *Bam*HI/*Xba*I, yielding plasmid pBBR1-MCS5::*eveD*. This construct was introduced by conjugation into the *S. fredii* HH103 *eveD*::Ω mutant, generating the complemented strain *S. fredii* HH103 *eveD*:: Ω (pBBR1-MCS5::*eveD*).

To analyse the expression profile of *eveD*, the promoter region and the first 12 bp of the coding sequence of *S. fredii* HH103 *eveD* gene were amplified using primers PeveD_XbaIF and PeveD_PstIR, yielding a 747-bp fragment. In parallel, primers PnopC_XbaIF and PnopC_PstIR were used to amplify a 383-bp fragment containing the *S. fredii* HH103 *nopC* promoter region and the first 9 bp of the coding sequence, to generate a reference transcriptional fusion for comparison. Both PCR products were digested with *Xba*I and *Pst*I and cloned upstream and in frame of *lacZ* into the pMP220 vector, generating pMP220::*_PeveD_*and pMP220::*_PnopC_*, respectively. These plasmids were introduced by conjugation into the *S. fredii* HH103 parental strain, as well as its mutant derivate strains affected in the *nodD1*, *nodD2*, *nolR*, *ttsI* and *syrM* genes.

For *Agrobacterium*-mediated transient expression, the *S. fredii* HH103 *eveD* ORF (∼5.1 kb) was amplified with primers eveD_attB1c_F and eveD_attB2c_R. The resulting product was cloned into pDONR207 (Invitrogen, USA) entry vector and then mobilised into pEarleyGate 101 binary vector (Earley *et al*., 2006) by the Gateway cloning system (Thermo Fisher Scientific, USA). Plasmids described above were introduced into *S. fredii* HH103 or *A. rhizogenes* K599 strains by triparental mating (Simon, 1984) using *E. coli* DH5α (pRK2013) as helper strain (Figurski and Helinski, 1979). In the case of *A. tumefaciens* GV3101, plasmid pEarleyGate 101::*eveD* was transformed by heat shock as described by Zhou *et al*. (2009). Standard recombinant DNA procedures were performed according to the general protocols (Sambrook *et al*., 1989).

### Bioinformatical analyses

Homologues of EveD (SFHH103_00272, 1687 aa) were retrieved from the NCBI non-redundant protein database by BLASTp (Altschul *et al*., 1997). Hits were retained as genuine homologues only when they aligned over at least 80 % of the query length with at least 45 % amino-acid identity; because the copy number of the tandem 136-residue repeats varies between strains, coverage was computed on the query rather than on the subject. These criteria excluded four partial or fragmentary entries and left 215 sequences. Redundancy was then removed by average-linkage clustering of the pairwise identity matrix at a 99 % threshold, retaining one representative per cluster; the EveD sequence of *S. fredii* HH103 was retained as its own leaf. The final dataset comprised 133 sequences representing the 215 homologues. It consists almost entirely of the order *Hyphomicrobiales* (*Rhizobiales*): 123 of the 133 leaves correspond to named species of this order, nine derive from metagenome-assembled genomes identified only to family or class level (*Rhizobiaceae*, *Alphaproteobacteria* or *Hyphomicrobiales* bacterium), and one, *Pseudomonas* sp. R2.Fl (MCL6707596.1, order *Pseudomonadales*), falls outside the order and was retained because it met the homology criteria. The full list of leaves, with accessions, organisms, lengths, clade assignment and identity to EveD, is given in Supplementary Dataset S1.

Sequences were aligned with FAMSA (Deorowicz *et al*., 2016), as implemented in pyfamsa v0.7.0. Poorly aligned regions, which correspond mainly to the variable repeat arrays, were removed by retaining only alignment columns with residue occupancy ≥ 80 %; this retains the columns aligned to the EveD sequence, yielding a final matrix of 1687 positions with 2.1 % gaps.

Evolutionary distances between all sequence pairs were estimated by maximum likelihood under the JTT amino-acid substitution model (Jones *et al*., 1992), optimising the branch length for each pair over the observed state-pair counts. The tree was then inferred by neighbour joining (Saitou and Nei, 1987). Branch support was obtained from 1000 nonparametric bootstrap replicates of alignment columns (Felsenstein, 1985), with the maximum-likelihood distances fully re-estimated for every replicate. The tree was rooted on the clade formed by the *Aquibium*, *Hoeflea*, *Rhizobium* sp. Leaf453 and unclassified *Rhizobiaceae* sequences (Clade 5), which is the most divergent lineage in the dataset (mean distance 0.62-0.84 substitutions per site to the other clades) and whose separation from the remaining sequences was recovered in 100 % of bootstrap replicates. Clades were delimited objectively as the maximal subtrees that combine bootstrap support ≥ 0.90 with a mean within-clade divergence ≤ 0.51 substitutions per site; this threshold corresponds to the upper limit of the within-clade divergence observed (0.13-0.51) and separates it from the divergence between clades (0.44-0.84). Seven clades were obtained, containing 125 of the 133 leaves; the remaining eight sequences, which sit on long isolated branches, were left unassigned. Support for merging clades was assessed by recording, in the same 1000 bootstrap replicates, how often the union of each pair of clades was recovered as a monophyletic group.

Phylogenetic analyses were run in Python 3.11 with pyfamsa 0.7.0 (FAMSA), Biopython 1.88 (Cock *et al*., 2009), DendroPy 5.1.3 (Moreno *et al*., 2024) and NumPy 2.4.6 (Harris *et al*., 2020); the JTT exchangeability matrix and equilibrium frequencies were taken from pyvolve (Spielman and Wilke, 2015). Sequence and annotation records were retrieved from NCBI with E-utilities. The tree is provided in Newick format with bootstrap values as internal node labels.

To analyse the sequence-based domain architecture of EveD, conserved domains were predicted and identified using the NCBI Conserved Domain Search (CD-Search) tool (Lu *et al*., 2020) against the Conserved Domain Database (CDD), and a consensus sequence logo was generated using Skylign (Wheeler *et al*., 2014) to visualise residue conservation across the EveD repeat units. The full-length EveD sequence was then submitted to SWISS-MODEL (Waterhouse *et al*., 2018) for comparative modelling on the AvrBs3/PthA structure, and the full-length three-dimensional structure was predicted independently of any template using AlphaFold (Abramson *et al*., 2024). Structural homology searches were performed on the resulting model using Foldseek (Van Kempen *et al*., 2024), which enables structure-based comparisons of protein models against structural databases through a structural alphabet encoding tertiary interactions. Secondary-structure content of the model was assigned with the P-SEA algorithm (Labesse *et al*., 1997) as implemented in Biotite 1.6.0 (Kunzmann and Hamacher, 2018), and per-residue pLDDT values were taken from the B-factor field of the model. Repeat modularity was assessed by least-squares superposition of consecutive 136-residue Cα segments onto the first unit, with half-period (68-residue) and quarter-period (34-residue) offsets as controls, and the maximum dimension and radius of gyration were computed from the Cα coordinates. To predict the EveD subcellular location, SignalP-6.0 (Teufel *et al*., 2022), DeepTMHMM (Hallgren *et al*., 2022, Preprint), DeepLoc 2.0 (Thumuluri *et al*., 2022), WoLF PSORT (Horton *et al*., 2007) and Localizer (Sperschneider *et al*., 2017) were used.

For the analysis of candidate secretion routes, the repertoire of protein secretion systems encoded by *S. fredii* HH103 was examined in the PGAP annotation of assembly GCF_048585425.2 and in the original annotation of assembly GCA_000283895.2, by searching the predicted proteomes for the diagnostic components of the type I (T1SS) to type VI (T6SS) systems and by inspecting the genomic neighbourhood of each candidate cluster; the resulting inventory was compared with that reported previously for this genome (Vinardell *et al*., 2015). RTX-type calcium-binding nonapeptides were searched across all 6187 predicted proteins of *S. fredii* HH103 using the GGxGxDxUx consensus (U = I, L, V, F, M, W or Y) and a relaxed G[GA]xGxD[^P]Ux variant. The amino-acid composition and net charge of the C-terminal 60 residues were computed for every protein of the proteome to provide a reference distribution.

Claude Opus 5 (Anthropic) was used to assist with the phylogenetic analysis, with the structural characterisation of the AlphaFold model (secondary-structure assignment, per-residue confidence, superposition of consecutive repeat units and molecular dimensions) and with the survey of candidate secretion routes. In each case the tool was used to write and refine the corresponding analysis code and to visualise the results. The remaining analyses described in this section, including comparative modelling, structural homology searches, domain and repeat-unit identification and subcellular localisation prediction, were carried out by the authors without such assistance.

### β-galactosidase assays

The *S. fredii* HH103 parental and derivate strains carrying the transcriptional fusions were pre-cultured in TY, YM or MM liquid media supplemented with the appropriate antibiotics. After incubation for 48 h under optimal growth conditions, 100 µl of each pre-culture was used to inoculate 3 ml of the corresponding fresh medium supplemented, where indicated, with genistein or soybean seed exudates. Cultures were grown to either the exponential (16 h; optical density at 600 nm [OD□□□] ∼ 0.6) or stationary (48 h; OD□□□_nm_ ∼ 1.2) growth phase. At the indicated time points, aliquots were collected for the determination of the β-galactosidase activity in permeabilised whole cells, as described by Zaat *et al*. (1987). β-galactosidase activity was expressed in Miller units according to the method described by Miller (1972). The same procedure was applied to the different *S. fredii* HH103 regulatory mutant backgrounds. In this case, β-galactosidase activity was determined at the exponential growth phase in MM liquid medium.

### RNA extraction and quantitative real-time PCR experiments

Total RNA was extracted as described by Vargas *et al*. (2011), with minor modifications. Briefly, bacteria grown in MM medium to the exponential growth phase were harvested, and frozen cell pellets were resuspended in Tri-Reagent (Invitrogen, USA; 2 ml per 25 ml of cell pellet). Samples were incubated at 65 °C for 20 min, and RNA was subsequently extracted and purified following the published protocol. Residual genomic DNA was removed by DNase treatment (Roche, Switzerland) at 37 °C for 1 h, followed by two sequential extractions with acid phenol:chloroform:isoamyl alcohol (25:24:1, v/v/v). RNA was finally precipitated overnight at -20 °C with ethanol and sodium acetate, washed with 70% ethanol, and resuspended in 50 µl of DEPC-treated water. RNA integrity was assessed by agarose gel electrophoresis and its concentration was assessed with a Nanodrop 1000 spectrophotometer (Thermo Scientific, USA) and a Qubit 2.0 Fluorometer (Invitrogen, USA). Genomic DNA-free RNA was reverse transcribed into cDNA using the PrimeScript RT reagent Kit with gDNA Eraser (Takara, Japan). Quantitative real-time PCR (qRT-PCR) was performed using a LightCycler 480 (Roche, Switzerland) with the following conditions: 95 °C for 10 min; 95 °C for 30 s; 55 °C for 30 s; 72 °C for 15 s; 40 cycles, followed by a melting curve profile from 60 to 95 °C to verify the specificity of the reaction. Expression levels were normalised against the *S. fredii* HH103 16S rRNA gene, which served as the internal reference. Three biological replicates were analysed, each with six technical replicates, and fold-changes were calculated using the ΔΔCt method (Pfaffl, 2001). The selected genes and primers are listed in Supplementary Table S2.

### Purification of S. fredii HH103 cellular and extracellular proteins

Cellular and secreted (extracellular) proteins from *S. fredii* HH103 cultures were extracted using the method described by Reyes Pérez *et al*. (2025), with minor modifications. Briefly, *S. fredii* HH103 strains were grown in liquid MM or YM supplemented with the appropriate antibiotics for 48 h at 28 °C with shaking. Subsequently, 1 ml of the grown culture was used to inoculate 40 ml of the corresponding medium in the presence or absence of 3.7 μM genistein, and cultures were incubated for 48 h under the same conditions. To obtain the cellular protein extracts devoid of secreted proteins, the optical density of each culture was adjusted to an OD_600nm_ of 1. Cultures were then centrifuged at 16.000 *g* for 1 min, the supernatant was discarded, and the resulting pellet was resuspended in 100 μL of sample loading buffer [62.5 mM Tris-HCl (pH 6.8), 2% (w/v) SDS, 10% (v/v) glycerol, 5% (w/v) β-mercaptoethanol, and 0.001% (w/v) bromophenol blue]. Extracellular proteins were isolated from 15 ml of culture by performing four consecutive centrifugation steps at 5.000 *g* for 20 min at 4 °C, discarding the pellet after each centrifugation to ensure complete removal of bacterial cells. Then, 1.8 ml of the resulting supernatant was transferred to a microcentrifuge tube containing 200 μl of 10% (v/v) trichloroacetic acid, mixed gently by pipetting and incubated overnight at 4 °C. The mixtures were then centrifuged at 16000 *g* for 30 min at 4 °C, and the supernatant was discarded. The protein pellet was washed with 1 ml of ice-cold 90% acetone and centrifuged again under the same conditions. Finally, the pellet was air-dried at room temperature until the acetone was completely evaporated and resuspended in sample loading buffer.

### Isolation and purification of S. fredii HH103 bEVs

*S. fredii* HH103 bEVs were purified following the protocol described by Ayala-García *et al*. (2024), with some modifications. Briefly, 10 ml of the *S. fredii* HH103 parental strain culture grown in MM at 28 °C for 72 h by shaking were used to inoculate in 1 l of the same medium in the presence or abscense of 3.7 μM genistein and incubated for 48 h under the same conditions. Then, cultures were centrifuged at 5000 *g* for 20 min at 4 °C. The resulting supernatant was filtered under vacuum using a LABOPORT® UN86 vacuum pump (KNF, Germany) fitted with 0.45 μm membrane vacuum filters (VWR, USA). The filtered supernatant was then concentrated to a final volume of 25 ml using a Vivaflow® 50R ultrafiltration system equipped with Vivaflow® 50R Hydrosart membranes (30,000 MWCO; Sartorius, France), according to the manufacturer’s instructions. The concentrated supernatant was subsequently subjected to ultracentrifugation for bEV purification. The concentrated supernatant was transferred to ultracentrifuge tubes and centrifuged at 100,000 *g* for 2 h at 4 °C using a Beckman Optima L-90K ultracentrifuge (GMI, USA) fitted with a 45 Ti rotor. Following ultracentrifugation, the supernatant was carefully removed from the pellet using a micropipette, yielding an bEV-containing pellet and an bEV-depleted supernatant. The pellet was resuspended in 300 μL HEPES buffer (Sigma-Aldrich, USA) and stored at 4 °C. Finally, proteins from each fraction were precipitated as described above up to the recovery of the cell-free supernatant, after which proteins were precipitated by adding a proportional volume of 10% (v/v) TCA. Proteins from the bEV fraction and the bEV-depleted supernatant were extracted following the procedure described above for extracellular protein extraction. When required, proteins from the concentrated supernatant were extracted using the same protocol. In all cases, the volume of 10% (v/v) TCA was adjusted proportionally to the sample volume.

### Quantification of rhizobial bEVs

bEV concentration and size distribution were determined as described by Ayala-García *et al*. (2024). Briefly, nanoparticle tracking analysis (NTA) was used based on light scattering using 1000X concentrated bEV samples. Measurements were performed with a NanoSight NS300 nanoparticle analyser (Malvern Panalytical, UK) equipped with a 488 nm monochromatic laser. Videos of 90 s were recorded at 23 °C and analysed using the manufacturer’s nanoparticle tracking analysis software to determine bEV concentration and particle size distribution according to the manufacturer’s instructions.

### Protein analysis of cellular, extracellular and bEV-contained fractions

For immunodetection, proteins were resolved on 4-20% (w/v) SDS-polyacrylamide gradient gels (Bio-Rad, USA) and total protein was visualised using a Gel Doc™ XR imaging system (Bio-Rad, USA) prior to protein transfer onto the membrane. Then, proteins were transferred onto nitrocellulose membranes (Bio-Rad) using a Mini Trans-Blot electrophoretic transfer cell (Bio-Rad). For transfer, the gel and membrane were positioned in direct contact and assembled between Whatman filter papers pre-equilibrated in chilled transfer buffer (48 mM Tris, 39 mM glycine, 1.3 mM SDS and 20% methanol). Proteins were transferred at 25 V and 1 A for 20 min. Then, membranes were blocked in TBS (0.2 M Tris, 1.3 M NaCl, pH 7.5) supplemented with 5% (w/v) skimmed milk for 1 h to prevent non-specific antibody binding, before being incubated overnight at 4 °C with a specific anti-EveD primary antibody (Davids Biotechnologie, Germany) diluted 1:100 in the same blocking solution. After washing three times with TBS-Tween buffer [TBS containing 1 mM Tween 20] for 5 min each at room temperature, membranes were incubated for 1 h at room temperature with an HRP-conjugated anti-rabbit secondary antibody (Cell Signaling Technology, USA). After incubation, membranes were washed three times with TBS-Tween buffer for 5 min each at room temperature. Immunoreactive bands were detected using Immobilon Forte HRP substrate (Merck, Germany) according to the manufacturer’s instructions and visualised with the LI-COR Odyssey FC 700/800/Chemi imaging system and analysed with Image Studio Lite software (LI-COR Biosciences, USA).

### Hairy Root Transformation in Soybean for Subcellular Localisation of EveD

To investigate the subcellular localisation of *S. fredii* HH103 EveD in root cells of its natural host plant, *G. max* cv. Williams, transient expression assays were performed by generating transgenic hairy roots following transformation with *A. rhizogenes* K599 (Supplementary Fig. S2). The procedure was based on the method described by Mohammadi-Dehcheshmeh *et al*. (2014), with minor modifications. *A. rhizogenes* K599 strains harbouring the corresponding plasmids were streaked onto LB agar plates supplemented with the appropriate antibiotics and incubated at 28°C for 24 h. Bacterial biomass was then collected with a sterile needle and used to inoculate soybean seedlings by direct injection into the hypocotyl approximately 1 cm below the cotyledons. Inoculation was performed on seedlings grown in vermiculite for approximately 1 week, when the cotyledons were fully expanded. The inoculation site was covered with sterile vermiculite, and plants were enclosed with a transparent plastic or glass cover to maintain high humidity. Plants were irrigated with nitrogen-supplemented Fåhraeus solution (Vincent, 1970), pH 6.8, and maintained under controlled environment conditions in a plant-growth chamber with a 16-h day, 8-h night cycle (light intensity of 500 µE x m−2 x s−1) and a relative humidity of 50%. Growth temperatures were set to 26 °C during the day and 18 °C during the night.

Three to four weeks after transformation, hairy roots expressing EveD-YFP fusion proteins were identified using a Zeiss AxioZoom V16 stereomicroscope (Zeiss, Germany). Transformed roots were mounted on glass slides in 10 mM MgCl□. When required, samples were stained with 4′,6-diamidino-2-phenylindole (DAPI) at a final concentration of 2 μg ml□¹ in 10 mM MgCl□. Coverslips were then applied and sealed prior to imaging with a Leica SPE confocal laser-scanning microscope (Leica Microsystems, Germany). Images were processed using LAS X Office software.

### Agrobacterium-mediated transient expression on Nicotiana benthamiana leaves and assessment of subcellular localisation of EveD

To investigate the specific subcellular localisation of EveD, transient expression assays were performed in *N. benthamiana* as previously described by Roden *et al*. (2004), with some modifications. *N. benthamiana* plants (Goodin *et al*., 2008) were grown in a growth chamber under controlled conditions (16□h at 26□°C in the light and 8□h at 18□°C in the dark, 70% humidity). For agroinfiltration, 100 μL of overnight cultures of *A. tumefaciens* GV3101 harbouring the corresponding binary vectors were inoculated onto 5 ml of LB medium supplemented with 40 μM acetosyringone (AS), 10 mM 2-(N-morpholino)ethanesulfonic acid (MES), and the appropriate antibiotics and incubated for 16 h at 28 °C with shaking. Then, cultures were harvested by centrifugation (3500 *g*, 5 min), washed with sterile 10 mM MgCl□, and resuspended in sterile 10 mM MgCl□ with 150 μM AS and 10 mM MES (pH 5.6). Bacterial suspensions were adjusted to an OD_600nm_ of 0.8 and incubated at 25 °C for 3 h without shaking before infiltration into the abaxial surface of leaves from 4-week-old *N. benthamiana* plants using a needleless syringe.

Subcellular localisation of EveD was analysed 48 h following transient expression of *A. tumefaciens* GV3101 carrying the pEarleyGate 101 construct expressing EveD fused in-frame to a C-terminal yellow fluorescent protein (YFP)-HA tag. When indicated, co-localisation analyses were performed by co-expressing EveD-YFP with fluorescent subcellular markers, including the monomeric red fluorescence protein (mRFP) fused in-frame with the endoplasmic reticulum marker HDEL (mRFP-HDEL; Schoberer *et al*., 2009); and the endosome marker FYVE fused in-frame with DsRed (Voigt *et al*., 2005). Alternatively, nuclei were stained with 4′,6-diamidine-2′-phenylindole dihydrochloride (DAPI, 1 mg ml□¹), while chloroplasts were identified by their autofluorescence. *A. tumefaciens* GV3101 derivative strains carrying pEarleyGate 104 and pEarleyGate 100 were used as positive and negative controls, respectively. Fluorescence signals were examined by confocal laser-scanning microscopy using a Leica SPE confocal microscope (Leica Microsystems, Germany). Z-stack images were acquired and processed with ImageJ software (Schneider *et al*., 2012), applying identical contrast and intensity settings to all images within each experiment.

### Plant tests

Nodulation assays to evaluate the symbiotic phenotype of the *S. fredii* HH103 *eveD* mutant in *G. max* cv. Williams 82, as well as nodulation assays with *G. soja* JL2 and CH2 lines were performed using a pot-based system. Soybean plants were grown individually in 800 ml pots containing expanded clay, with ten pots per treatment, each containing a single plant. Pots were arranged in separate trays according to treatment to prevent cross-contamination, with each tray containing 6 L of Fåhraeus (Vincent, 1970), pH 6.8, The substrate, pots and nutrient solution were sterilised by autoclaving at 121 °C for 20 min, whereas the trays were disinfected with 98% (v/v) ethanol. The pots were then filled with the sterile substrate, and a single seed was sown in each pot. Each seed was inoculated with 1 ml of a bacterial culture grown in YM medium to an OD_600nm_ of 0.6. All procedures, including the addition of sterile nutrient solution to the trays, were carried out under aseptic conditions in a laminar flow hood. Plants were grown under a controlled environment chamber as detailed above. Nodulation parameters, including the number of nodules, pseudonodules (if applicable), nodule fresh mass, and shoot dry mass, were evaluated after 42 days. The shoots were dried at 70 °C for 48 h and then weighed.

Coinoculation experiments (competitiveness) on *G. max* cv. Williams 82 were performed using the parental strain *S. fredii* HH103 and its mutant derivative *S. fredii* HH103 *eveD*::Ω, following the methodology described by Reyes-Pérez *et al*. (2025), with minor modifications (Supplementary Fig. S3). Bacterial strains were grown in YM medium to an OD_600nm_ of 1.0. Germinated soybean seeds were inoculated with 1 ml of a bacterial mixture at three distinct ratios: 1:1, 10:1, and 1:10. Plants were cultivated for 30 days in a growth chamber under the environmental conditions described above. To determine nodule occupancy, 100 nodules per treatment were surface-sterilised by immersion in 6% (v/v) sodium hypochlorite for 6 min, followed by six consecutive washes with sterile distilled water. Sterilisation efficacy was verified by rolling the treated nodules onto antibiotic-free TY agar plates to confirm the absence of superficial growth. Individual nodules were crushed with sterile forceps, and the rhizobia-containing exudate was plated onto TY supplemented with Rif, and TY supplemented with Rif and Spc, being the plates incubated at 28 °C for 48 h. Phenotypic identification was based on antibiotic resistance profiles (the parental strain is resistant only to Rif, whereas the mutant strain exhibits resistance to both Rif and Spc). Nodules displaying growth exclusively on TY+Rif plates were scored as occupied solely by the parental strain. For nodules yielding bacterial growth on both selective media, the presence of mixed occupancy or exclusive mutant colonisation was determined by resuspending the biomass recovered from the initial TY+Rif plate in 1 ml of liquid TY medium. Serial dilutions were subsequently plated onto TY+Rif agar, and 100 randomly selected colonies from plates containing 100-1000 colonies were replica-plated onto TY+Rif and TY+Rif+Spc media. The proportion of each strain was then calculated, allowing the identification of nodules occupied by a single strain or co-occupied by both strains.

### Statistical analysis

The statistical tests performed in this work are indicated in the figure legends and were done using Prism 8 (GraphPad, La Jolla, CA, USA).

## Results

### Identification and bioinformatic characterisation of EveD, a large rhizobial repeat-containing protein

This study originated in a computational screen of the *S. fredii* HH103 genome with T346Hunter (Martínez-García *et al*., 2015), a tool developed to detect components of the T3SS, type IV (T4SS) and T6SS secretion systems and later integrated into the Plant Interaction Factors Resource (PIFAR) database (Martínez-García *et al*., 2016). The screen flagged *SFHH103_00272*, hereafter *eveD*, as a candidate T3E on the basis of sequence similarity to the TAL effector gene *avrBs3* of *X. campestris* (Bonas *et al*., 1989; Boch and Bonas, 2010; Deng *et al*., 2012). To place EveD (1687 aa) among related bacterial proteins, we built a phylogeny from full-length homologues of rhizobia and related genera, almost all of them belonging to the order *Hyphomicrobiales* (Fig. 1). Of the 219 sequences retrieved, 215 met the homology criteria and were grouped at 99 % identity into 133 representatives (Supplementary Dataset S1). The tree resolved seven well-supported clades that together contained 125 of the 133 representatives, whereas the branching order among the clades remained unresolved, indicating that the family diversified early and rapidly. EveD falls within Clade 4 (n = 15), which comprises a tight core of ten *Sinorhizobium* and *Ensifer* proteins of near-uniform length (nine of 1687 aa and one of 1719 aa; 90.6-100 % identity to EveD) together with a more distant subgroup of five *Rhizobium* and *Rhizobiaceae* sequences (69.9-76.8 % identity; 1415-1823 aa). The remaining clades are considerably more divergent from EveD (47-61 % identity) and differ in protein length, Clade 1 being the largest (n = 43; median length 2435 aa).

**Fig. 1.**
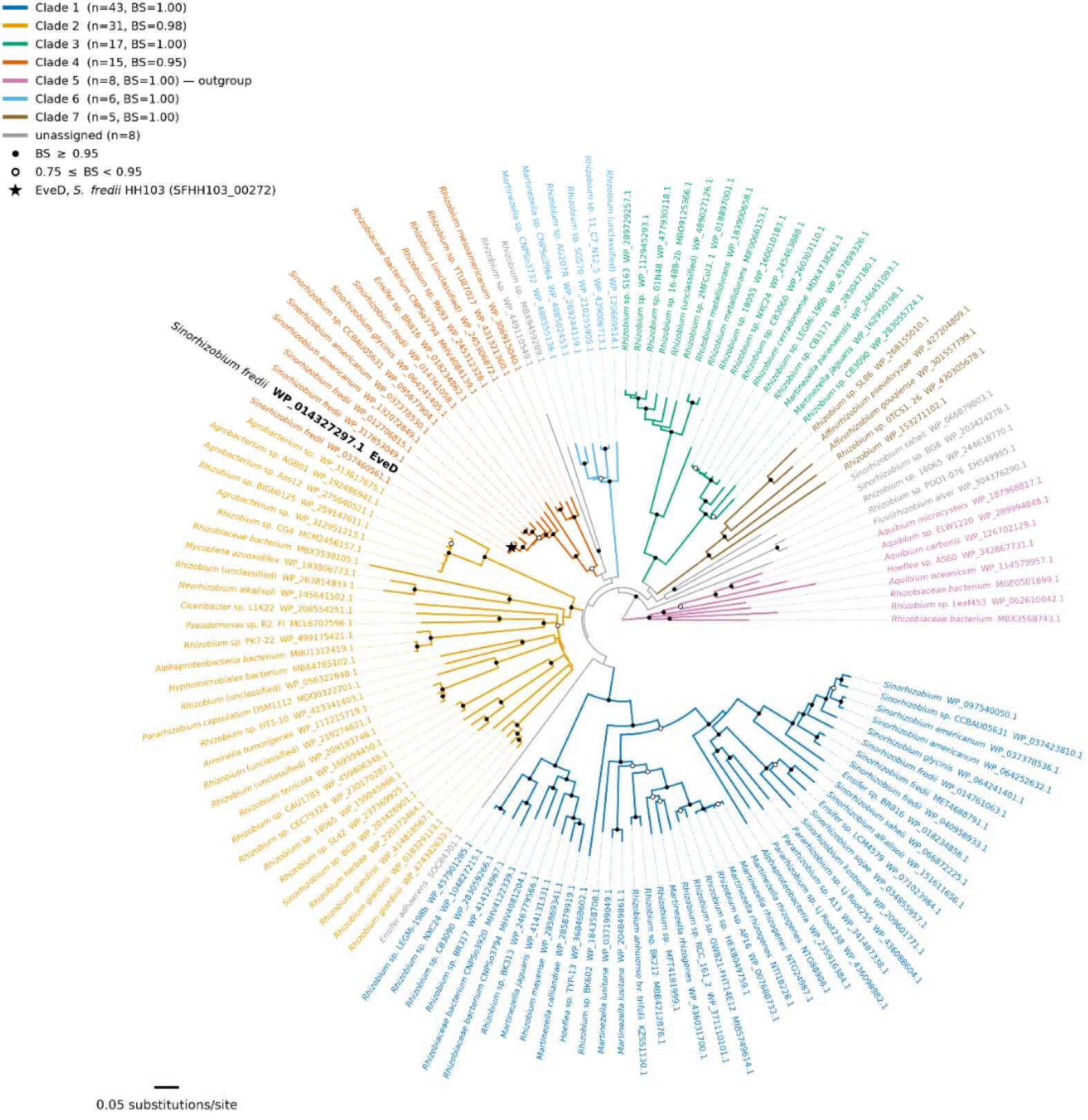
Phylogeny of the EveD-like protein family in *Sinorhizobium*, *Rhizobium* and related genera. Homologues of EveD were retrieved by BLASTp and filtered to retain full-length relatives (≥45 % identity and ≥80 % query coverage), leaving 215 sequences that were clustered at 99 % identity into 133 tree leaves (the accessions collapsed into each leaf are listed in Supplementary Dataset S1). Sequences were aligned with FAMSA and the alignment trimmed to the 1687 columns with ≥80 % residue occupancy (2.1 % gaps). Pairwise distances were estimated by maximum likelihood under the JTT model and the tree was inferred by neighbourjoining method; node support is from 1000 bootstrap replicates in which distances were re-estimated from the resampled columns (filled circles, BS ≥ 0.95; open circles, 0.75 ≤ BS < 0.95; 100 of the 130 internal branches reach BS ≥ 0.75). The tree is rooted on Clade 5 (*Aquibium*, *Hoeflea* and unclassified *Rhizobiaceae*), the most divergent lineage recovered and the only split supported in every bootstrap replicate (BS = 1.000). Colours mark the seven clades delimited as maximal subtrees with BS ≥ 0.90 and mean within-clade divergence ≤ 0.51 substitutions per site. Together they contain 125 of the 133 leaves, and the eight leaves outside any such clade are grey. Because the branching order among clades is unresolved (BS ≤ 0.45 for every pairwise union of clades), their circular arrangement carries no phylogenetic meaning. The black star marks EveD of *S. fredii* HH103 (*SFHH103_00272*), which falls within Clade 4 (BS = 0.95). Of the 133 leaves, 123 are named species of the order *Hyphomicrobiales*, nine derive from metagenome-assembled genomes assigned only to family or class level, and one (*Pseudomonas* sp. R2.Fl, Clade 2) lies outside the order. Scale bar, 0.05 substitutions per site.

The EveD sequence contains an extended array of tandemly arranged internal repeats: eleven conserved units of 136 amino acids spanning the central region of the protein, with a mean pairwise identity of 80.3 % and a strongly conserved consensus (Supplementary Fig. S4). In line with the original screen, conserved-domain analysis with the NCBI CDD returned similarity to FhaB-like proteins and to the AvrBs3 family of *Xanthomonas*, and a comparative model built with SWISS-MODEL on the AvrBs3/PthA structure (PDB 3UGM) aligned residues 640-1314 of EveD at 33.5 % identity, corresponding to part of the repeat region. The limited coverage and low confidence of that model, however, did not support classifying EveD as a canonical TAL effector.

We therefore examined the structure directly, using an AlphaFold model of the full-length protein. Structural and profile-based searches with Foldseek retrieved no canonical TAL effector, returning instead proteins with extended repetitive regions, such as ice-nucleation-like proteins (AF-A0A5Q0C5Y9-F1-model_v6; *Rhizobium grahamii*; Prob. 1.00; Seq. Id. 4.6 %; E-value 0.00e+0; query positions 1-1271) and haem-utilisation proteins (AF-A0A2U1IKW8-F1-model_v6; *Pseudomonas* sp. B1(2018); Prob. 1.00; Seq. Id. 43 %; E-value 7.64e-86; query positions 181-1425). Because the characterised members of all of these families adopt β-solenoid architectures, we quantified the secondary-structure content of the model: it is 70.5 % α-helical and contains no β-strand, and is built from 197 short helices (median length 6 residues, longest 13), an organisation characteristic of α-solenoids, not of the long continuous helices of a coiled coil. Per-residue confidence is high and uniform along the molecule (mean pLDDT 92.6; 88.0 % of residues above 90; none below 50), indicating that the low global score (pTM = 0.44) reflects uncertainty in the relative arrangement of the modules rather than in their fold. Consecutive 136-residue units superposed onto the first with root-mean-square deviations of 0.27-0.81 Å (median 0.48 Å), against 3.5 and 4.4 Å for controls offset by 68 and by 34 residues, the length of a TAL effector repeat. The repeat is thus a genuine α-helical module, incompatible with the β-solenoid architecture of the FhaB, ice-nucleation and RTX families and with the TAL effector fold. The modelled molecule is extended but curved (maximum dimension 271 Å, radius of gyration 77 Å).

EveD has no N-terminal Sec or Tat signal and no predicted transmembrane segment, indicating that it is a soluble protein, and subcellular localisation predictors place it extracellularly in bacteria and, in eukaryotic cells, in the cytoplasm, in agreement with the absence of a nuclear localisation signal or of any other recognised targeting motif. Because EveD lacks a signal peptide, we examined whether any other secretion system encoded by *S. fredii* HH103 could account for its export. The genome encodes two complete T1SS (*prtDE*, *SFHH103_03015-03016*, on the chromosome and *prsDE*, *SFHH103_06228-06229*, on plasmid pSfHH103e), the core components of a T2SS (a GspD secretin together with GspE, GspF, GspG, GspK, GspM and a prepilin peptidase), the symbiotic T3SS, and three plasmid-borne *virB* clusters, two of them complete and one truncated, in agreement with the inventory reported for this genome (Vinardell *et al*., 2015). By contrast, no gene encoding a classical autotransporter, a TpsB/FhaC-family transporter, a trimeric autotransporter adhesin or an intimin/invasin homologue was detected, the only related gene being the autotransporter assembly factor TamA (*SFHH103_03105*), and no component of a T6SS was found, in agreement with that same analysis and in contrast to the system described in *S. fredii* USDA257 (Reyes-Pérez *et al*., 2025). EveD contains no RTX-type calcium-binding nonapeptide, whereas the same search recovered between 10 and 29 copies in each of the eight *S. fredii* HH103 proteins annotated as haemolysin-type calcium-binding or RTX-like proteins, with which EveD shares no detectable sequence similarity; accordingly, EveD is not among the 18 proteins previously proposed as putative T1SS substrates in this genome (Vinardell *et al*., 2015). T2SS substrates must first reach the periplasm through the Sec or Tat pathways, which EveD cannot use, and T4SS secretion delivers substrates by contact-dependent translocation rather than releasing them into the cell-free medium. The evidence for and against each route is summarised in Supplementary Table S4.

Finally, *eveD* is transcribed independently of its neighbours: it lies on the strand opposite to both flanking genes, diverging from *SFHH103_00273* across a 731 bp intergenic region at its 5′ end and converging with *SFHH103_00271* across 356 bp at its 3′ end. *eveD* therefore constitutes a monocistronic transcription unit, so that the Ω interposon used to inactivate it cannot exert polar effects on co-transcribed downstream genes.

### Expression of the S. fredii HH103 eveD gene is independent of the canonical NodD-TtsI regulatory cascade

To determine the conditions regulating *eveD* expression, transcriptional activity was analysed using a P*_eveD_*::*lacZ* fusion in the parental *S. fredii* HH103 strain. The expression profile of *eveD* was compared with the flavonoid-inducible T3E gene *nopC* using a P*_nopC_*::*lacZ* fusion. The *lacZ*::promoterless vector pMP220 was included as a negative control. As a preliminary assessment to identify the most suitable culture conditions for subsequent experiments, β-galactosidase activity was determined in bacterial cells grown in standard (YM) and minimal (MM) media during exponential and stationary phases, in the absence or presence of genistein or soybean seed exudates.

As shown in Fig. 2, *eveD* displayed a broadly similar expression profile in YM and MM, with no marked differences attributable to the culture medium and/or the presence of plant-derived signals. In contrast, *nopC* exhibited the characteristic expression pattern of a T3SS-associated gene, responding strongly to both inducing signals under all the conditions tested (between 11.4-fold and 30.4-fold). Besides, its expression did not differ significantly between exponential and stationary phases in either medium.

**Fig. 2.**
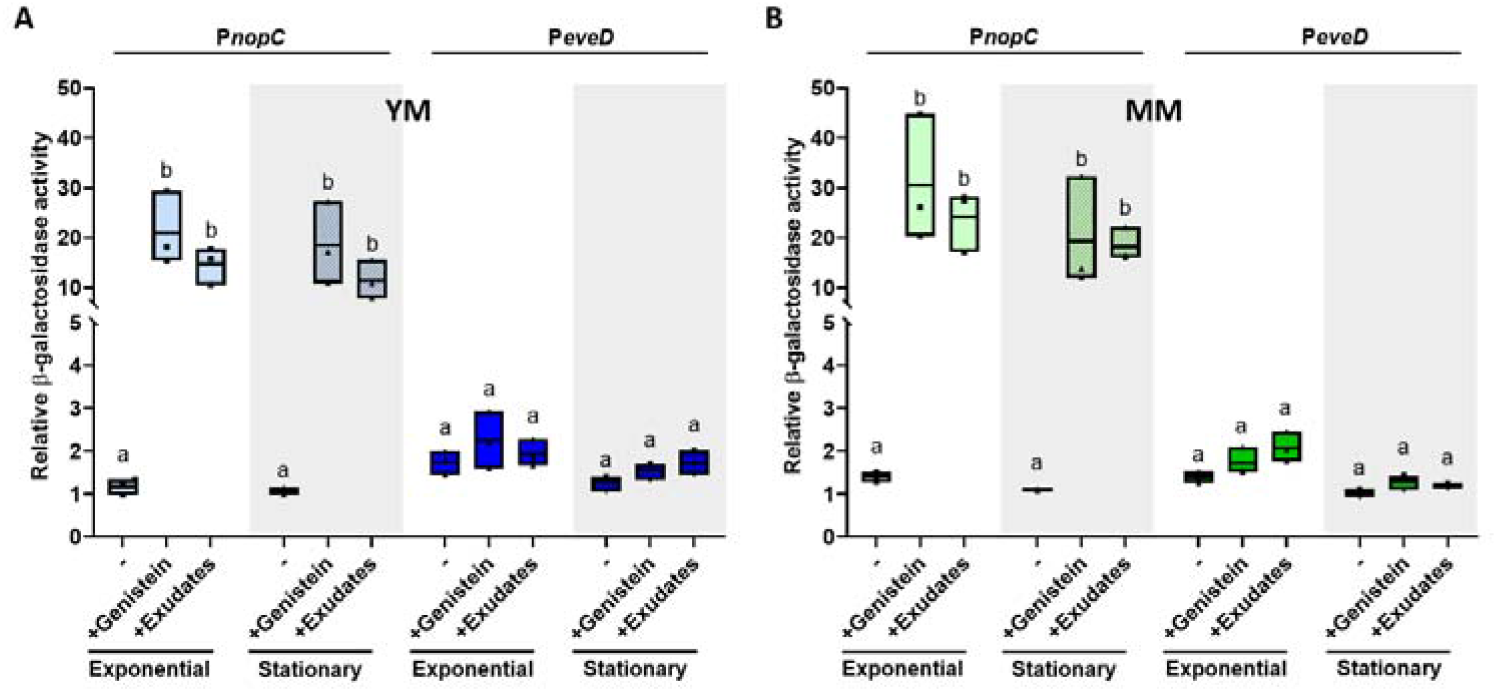
Activation pattern of the *S. fredii* HH103 *nopC* and *eveD* promoters under different culture conditions. Fold-change values of the β-galactosidase activity of HH103 carrying a pMP220 plasmid containing the *nopC* and *eveD* promoter regions fused to the *lacZ* gene with respect to those values obtained by the strain carrying the empty pMP220 plasmid. The tested conditions included YM (**A**, blue) and MM (**B**, green) media in the presence (+) or absence (-) of genistein or seed exudates evaluated at the exponential or stationary (striped) phases of the growth curve. Values are the mean ± standard deviation of three biological replicates, each assayed in three technical replicates. For each panel, groups labelled with the same letter are not significantly different at α = 5% (One-Way ANOVA with multiple comparisons, *P* < 0.05). At least three independent experiments were performed for each condition.

Sequence analysis of the *eveD* promoter region revealed a putative *tts* box and a SyrM box-like sequence (Supplementary Fig. S5). However, both motifs displayed deviations from the consensus sequences described for functional regulatory elements, suggesting that establishing their contribution to *eveD* regulation would require experimental validation. Therefore, β-galactosidase assays were performed in a set of mutant backgrounds affected in key symbiotic regulators to determine whether *eveD* expression depends on the NodD1/TtsI regulatory cascade or on other transcriptional regulators involved in symbiotic gene expression, including NodD2, NolR, and SyrM. The expression pattern of *nopC* was analysed in parallel as a control for a canonical T3SS-associated gene. All strains were grown in MM medium and harvested during the exponential phase.

As expected, *nopC* expression was strongly affected in the absence of the NodD1/TtsI regulatory cascade. In both the *nodD1* and *ttsI* mutant backgrounds, seed exudate-dependent induction was abolished compared with the parental strain (11.4-fold). In contrast, *nopC* expression was not affected in the *syrM* background (9.1-fold), whereas a slight but significant increase in expression was observed in the *nodD2* and *nolR* mutant backgrounds (18.4-fold and 23.0-fold, respectively). These results confirm that *nopC* activation under these conditions mainly depends on the NodD1/TtsI pathway and is repressed in the presence of NodD2 or NolR. Unlike *nopC*, *eveD* expression was not affected by the absence of NodD1 or TtsI regulators. The *nodD1* and *ttsI* mutants showed expression levels and induction patterns comparable to those of the parental strain, indicating that the *tts* box-like sequence present in the *eveD* promoter is not functional, at least in the tested conditions. Besides, *eveD* expression was not altered in the *nodD2* or *nolR* mutant backgrounds, showing expression levels comparable to those observed in the parental strain. However, a modest but statistically significant reduction in *eveD* expression was observed in the absence of SyrM independently of the presence of plant-derived signals (Fig. 3).

**Fig. 3.**
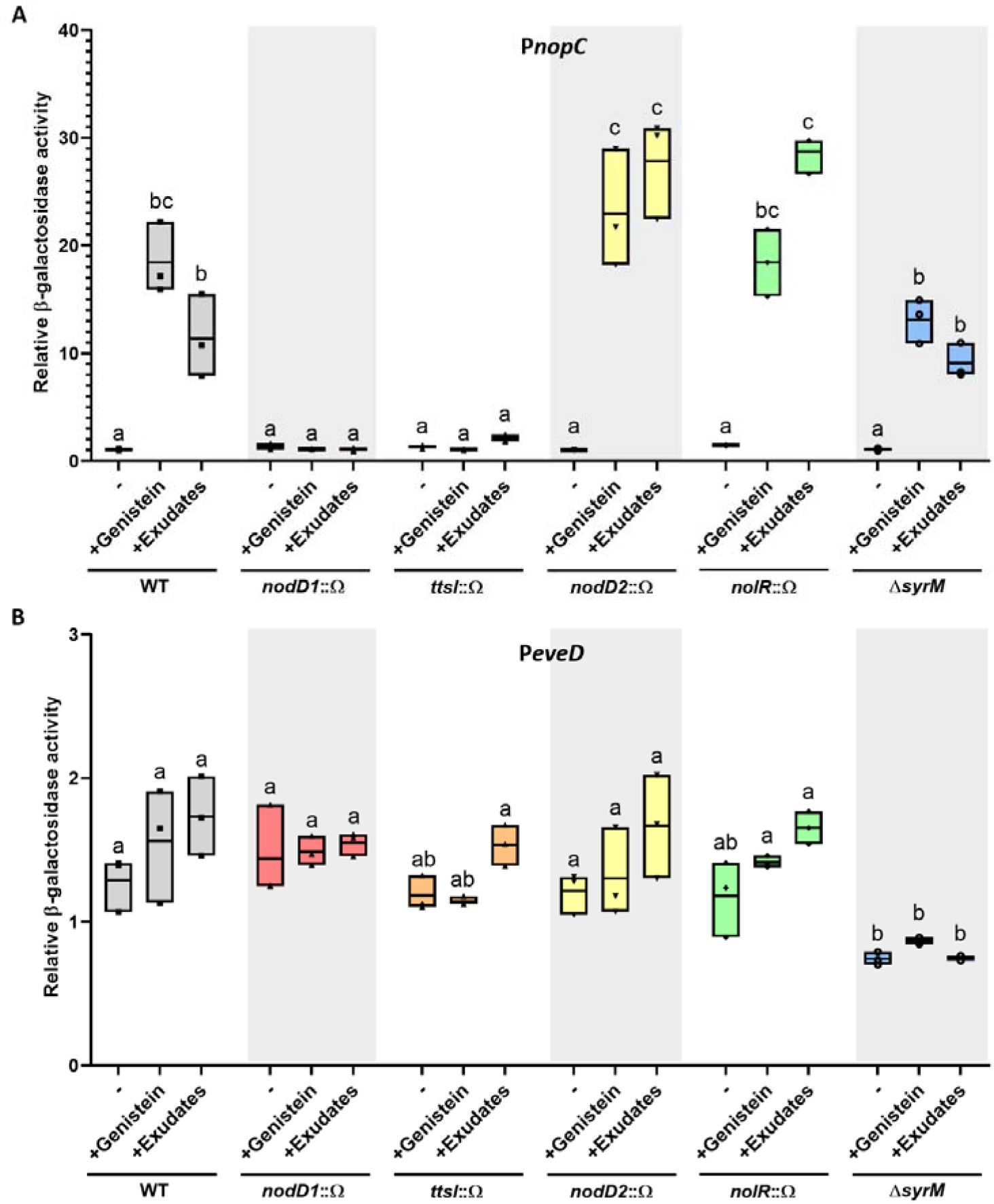
Activation pattern of *nopC* and *eveD* promoters in different *S. fredii* HH103 genetic backgrounds. Fold-change values of the β-galactosidase activity of HH103 carrying a pMP220 plasmid containing the *nopC* (**A**) and *eveD* (**B**) promoter regions fused to the *lacZ* gene with respect to those values obtained by the strain carrying the empty pMP220 plasmid. Wild-type (grey), *nodD1* (red), *ttsI* (orange), *nodD2* (yellow), *nolR* (green) and *syrM* (blue) strains were grown in MM medium in the presence or absence of genistein or seed exudates. Values are the mean ± standard deviation of three biological replicates, each assayed in three technical replicates. For each genetic background, groups labeled with the same letter are not significantly different at α = 5% (One-Way ANOVA with multiple comparisons, *P* < 0.05). At least three independent experiments were performed for each condition.

To further validate the regulatory pattern observed in the β-galactosidase assays, qRT-PCR analyses were performed to determine the expression levels of *eveD* and *nopC* in the different regulatory mutant backgrounds grown under non-induced conditions or in the presence of genistein. Based on the β-galactosidase assays, genistein was selected as the inducing condition for subsequent experiments as it produced a regulatory pattern comparable to that observed with soybean seed exudates while providing a defined and reproducible inducing condition. Thus, subsequent analyses focused on assessing the contribution of the different symbiotic regulators rather than comparing inducing signals.

The transcriptional profiles obtained by qRT-PCR (Supplementary Fig. S6) were consistent with the overall trends observed using the *lacZ* reporter fusions. In agreement with the reporter assays, *nopC* expression was strongly dependent on the activation by the NodD1/TtsI regulatory cascade. Besides, the *nodD2* and *nolR* mutant backgrounds showed a significant increase in transcript levels compared with the parental strain. Conversely, *eveD* transcript levels were not significantly altered in the majority of mutant backgrounds, except the *syrM* mutant, with a subtle but significant reduction in *eveD* expression compared with the parental strain.

Together, these results demonstrate that, despite the presence of imperfect *tts* and SyrM box-like sequences in its promoter region, *eveD* does not follow the canonical NodD1/TtsI-dependent regulatory pattern characteristic of T3SS-associated genes. Instead, *eveD* expression is partially dependent on the SyrM symbiotic regulator. To establish whether this relation depends on the presence of the SyrM box-like sequence located upsteram of *eveD* requires further investigation.

### EveD Secretion is Independent of a Functional T3SS and is Mediated by Bacterial Extracellular Vesicles

The secretion pattern of EveD was analysed by Western blot using extracellular (secretion) and whole-cell protein fractions from *S. fredii* HH103 cultures grown in MM medium either in the absence or in the presence of genistein. To assess whether EveD follows a canonical T3SS-dependent secretion pathway, its secretion pattern in different genetic backgrounds was compared with that of the rhizobial T3SS effector NopP using specific anti-NopP and anti-EveD antibodies (Fig. 4). As expected, NopP was detected in the extracellular fraction of the parental strain only under genistein-induced conditions, whereas its secretion was abolished in the *rhcJ* mutant, resulting in slight protein accumulation in whole-cell extracts. Similarly, NopP was not detected in the *ttsI* mutant, confirming its dependence on the NodD1/TtsI-regulated T3SS pathway.

**Fig. 4.**
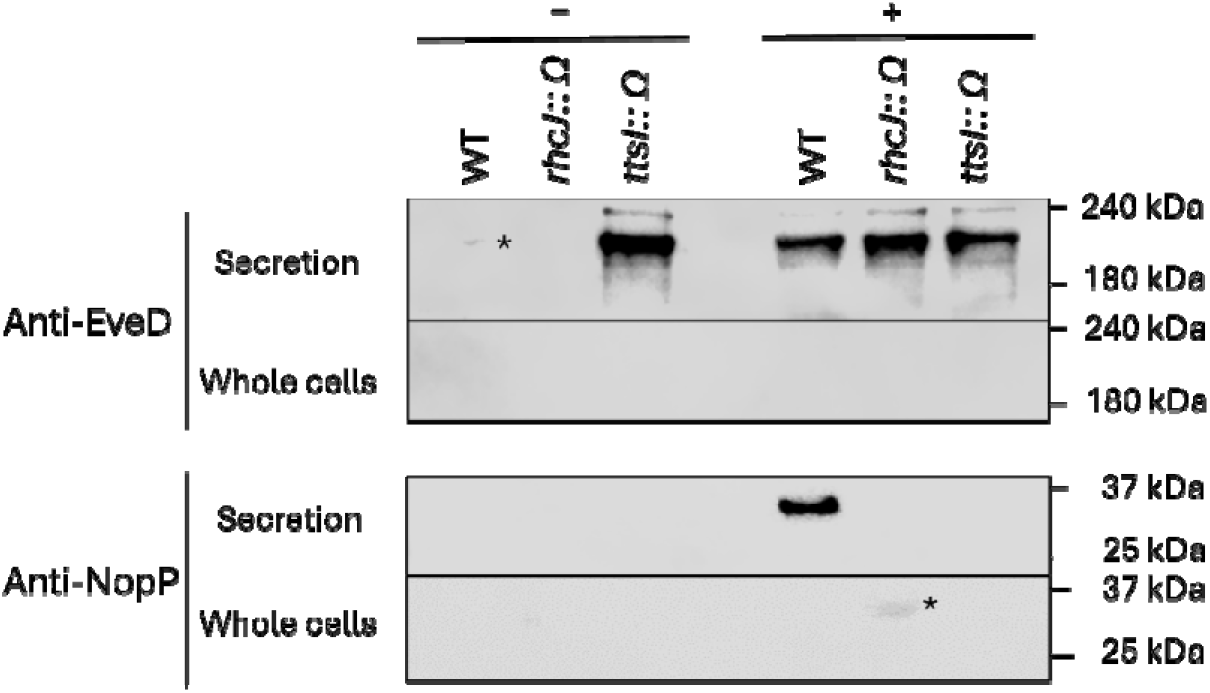
Secretion of the *S. fredii* HH103 EveD protein. Production and secretion of EveD and NopP in the wild-type strain and in the *rhcJ* and *ttsI* mutant derivatives grown in MM medium in the presence (+) or absence (-) of genistein. Proteins were detected by Western blot analysis using specific anti-EveD and anti-NopP antibodies. Specific EveD and NopP bands are marked with an asterisk, where necessary. The “Whole cells” panels represents the intracellular protein fraction, whereas “Secretion” panels displays the extracellular protein fraction. The position of the molecular size marker (in kDa) is indicated. The image is representative of three independent experiments.

In contrast, EveD was recovered from the extracellular fraction of both the *rhcJ* and the *ttsI* mutants in the presence of genistein, demonstrating that its export does not require a functional T3SS. The three backgrounds did not, however, behave identically under non-inducing conditions. In the parental strain extracellular EveD was barely detectable in the absence of genistein, and in the *rhcJ* mutant it was not detected at all, whereas in the *ttsI* mutant EveD accumulated extracellularly without the flavonoid to at least the same extent as in induced cultures. Thus, assembly of the secretion apparatus is dispensable for EveD export, but the amount of extracellular EveD under non-inducing conditions depends on the genetic background, being markedly higher in the absence of TtsI. Besides, under the conditions tested, EveD was not detected in whole-cell extracts of any of the strains analysed (Fig. 4).

Given that EveD secretion was independent of a functional T3SS, we investigated whether this protein could be exported through an alternative secretion mechanism. Since bEVs represent an emerging protein export pathway described in numerous bacterial species, including *S. fredii* HH103, we analysed the potential association of EveD with bEVs produced by this strain. Thus, bEVs were isolated and purified from cultures of the parental strain *S. fredii* HH103 grown in MM medium in the absence or presence of genistein. As we previously reported in Moreno-de Castro *et al*. (2026), particle concentrations increased in the presence of the inducing flavonoid (Supplementary Fig. S7). Protein samples from the different fractions obtained during the purification process were analysed by Western blot, including the initial cellular fraction (CF), filtered supernatant (S), concentrated supernatant (CS), bEV-free supernatant (S-EV) and bEV-enriched fractions (EV). Immunodetection was performed using a specific anti-EveD antibody and antibodies against the T3Es NopP and NopC, in order to compare the localization of EveD with that of known T3Es across bEV-associated and soluble fractions.

The results showed that NopP and NopC were detected exclusively in the presence of genistein. Interestingly, both T3Es were detected in extracellular fractions, including the bEV-free supernatant as well as in the bEV-enriched fraction, suggesting that, NopP and NopC may also be released through bEVs in *S. fredii* HH103 in addition to their canonical secretion via the T3SS. In contrast, EveD was detected in the filtered and concentrated extracellular fractions and in the bEV-enriched fraction under both non-induced and induced conditions but was absent from the bEV-free supernatantThe protein distribution pattern indicates that extracellular EveD is predominantly associated with bEV-containing fractions, supporting a secretion mechanism linked to bEVs rather than release as a soluble extracellular protein (Fig. 5; Supplementary Fig. S8).

**Fig. 5.**
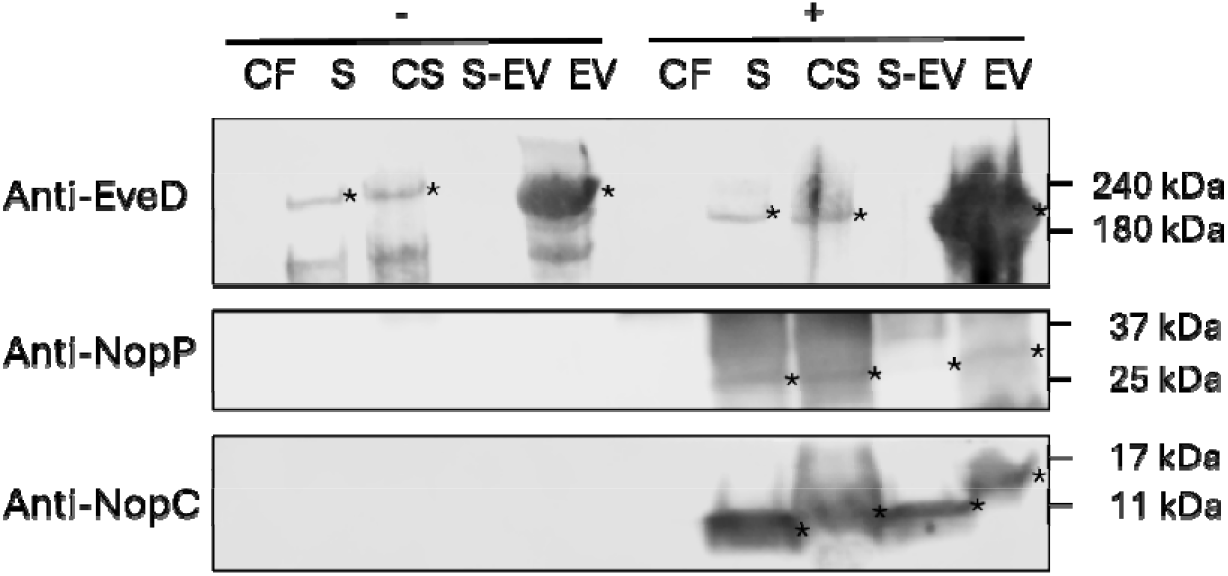
Presence of the *S. fredii* HH103 EveD protein in different cellular fractions. Production and secretion of EveD, NopP and NopC (marked with *) in the wild-type strain grown in MM medium in the presence (+) or absence (-) of genistein. Proteins were detected by Western blot analysis using specific anti-EveD, anti-NopP and anti-NopC antibodies. CF: cellular fraction, S: filtered supernatant, CS: concentrated supernatant, S-EV: EV-free supernatant, and bEV: bEV-enriched fractions. The position of the molecular size marker (in kDa) is indicated. The image is representative of three independent experiments.

Together, these findings support EveD as a non-canonical secreted protein whose extracellular release is closely associated with bEVs, consistent with an bEV-mediated delivery mechanism.

### EveD Localises Predominantly to the Plant Cytoplasm and Associates with the Endoplasmic Reticulum

bEVs have been shown to interact with plant cells (Tran *et al*., 2022; Bahar *et al*., 2016). Considering that *in silico* analyses indicate that EveD is a soluble protein lacking apparent transmembrane domains or other features associated with membrane localisation, and lacks predicted secretion signals compatible with conventional protein export pathways, we hypothesised that, following bEV-mediated delivery, EveD could be released into the plant cell cytoplasm.

To investigate this possibility, the subcellular localisation of EveD was analysed in cells of the natural host of *S. fredii* HH103, *G. max* cv. Williams. Transient expression assays were performed in soybean roots using the *A. rhizogenes* K599-mediated hairy root transformation system. This approach allows the generation of transgenic roots expressing proteins of interest in root cells. The construct pEarleyGate 101::*eveD*, which synthesizes EveD fused to the yellow fluorescent protein (YFP), was used to analyse EveD localisation by confocal microscopy. *G. max* cv. Williams roots were inoculated with *A. rhizogenes* K599 carrying pEarleyGate 101::*eveD* or, as control, pEarleyGate 104, which produces free YFP. Four weeks after inoculation, fluorescent roots were selected and analysed by confocal microscopy.

Unlike roots expressing free YFP, in which fluorescence was diffusely distributed throughout both the cytoplasm and nucleus of plant cells, EveD-YFP fluorescence showed a predominantly cytoplasmic localisation, showing occasional detection in nuclei (Fig. 6).

**Fig. 6.**
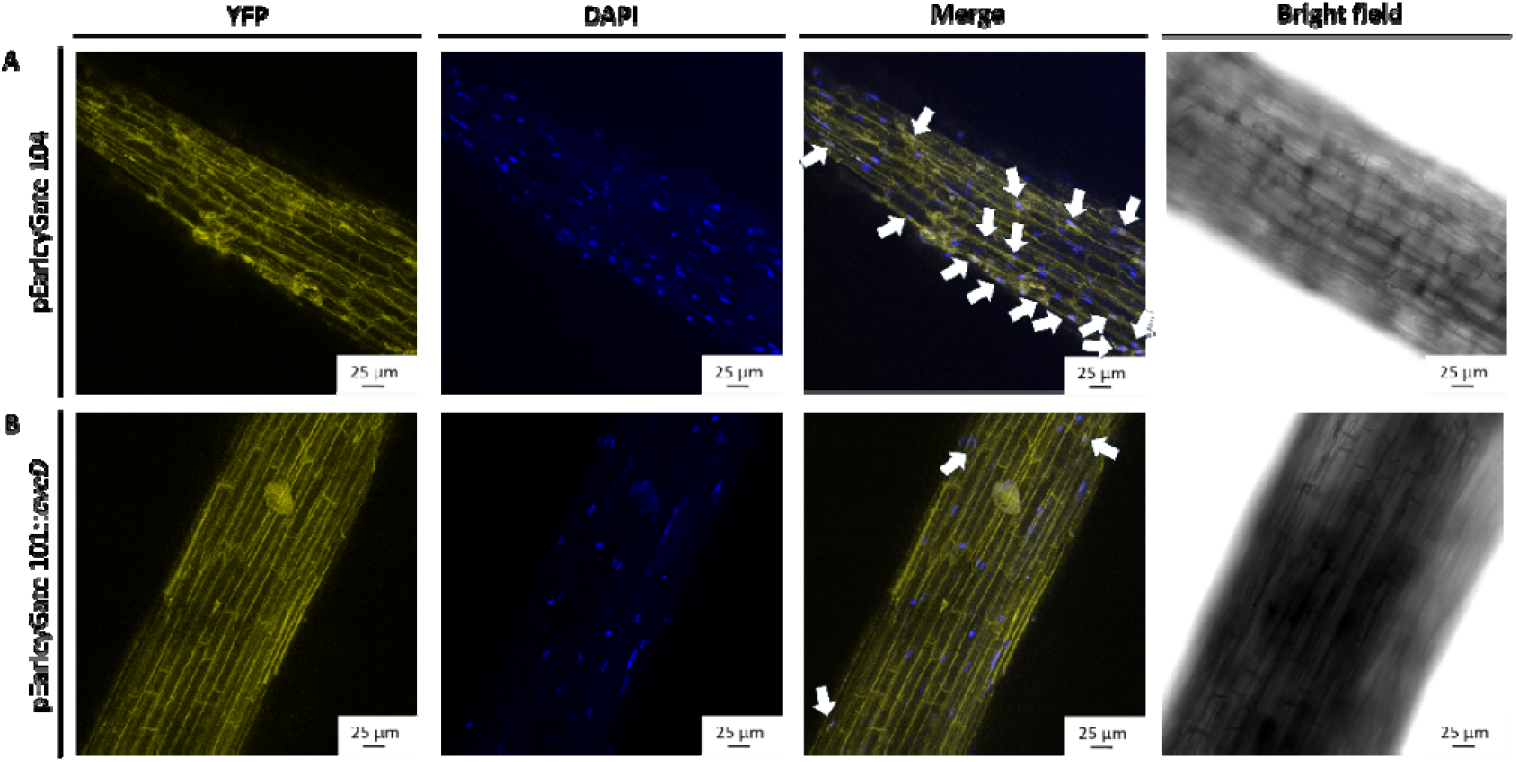
Subcellular localisation of *EveD* in *G. max* cv Williams root cells. Confocal images of soybean roots three weeks after agroinfiltration with *A. rhizogenes* K599. Nuclei are shown in blue (DAPI staining). Plasmid pEarleyGate 104 was used as a positive control for YFP expression (**A**). EveD localisation is shown in yellow (**B**, pEarleyGate 101::*eveD*). Scale bars indicate 25 µm. Arrows indicate nuclear localisation of YFP and EveD. Subcellular localisation of EveD was assessed in hairy root tissue derived from two independent transformation events.

Because the use of fluorescent compartment-targeting markers is technically limited in soybean roots, *N. benthamiana* leaves were used as a heterologous system to further define the subcellular distribution of EveD. These studies were performed by agroinfiltrating *A. tumefaciens* GV3101 carrying the plasmids named above. After 48 hours of agroinfiltration, infiltrated leaves were stained with DAPI and analysed by confocal microscopy.

Epidermal cells producing free YFP showed a diffuse fluorescence pattern throughout most cells (Fig. 7A). In contrast, EveD-YFP fluorescence was detected in a limited number of cells and displayed both cytoplasmic and nuclear localisation. Within the cytoplasm, EveD-YFP showed a homogeneous reticular distribution pattern (Fig. 7B), suggesting association with intracellular structures involved in cellular organisation or trafficking. To determine the cellular structures associated with this reticular distribution, co-infiltration assays were performed using *A. tumefaciens* GV3101 carrying pEarleyGate 101::*eveD* and a construct encoding the endoplasmic reticulum marker mRFP-HDEL. Confocal microscopy revealed colocalisation of EveD-YFP with mRFP-HDEL, indicating an association with ER (Fig. 7D). Since this structure participates in intracellular trafficking processes and membrane dynamics (Aniento *et al*., 2022), the possible association of EveD with endosomal compartments was subsequently investigated. Co-infiltration assays were performed using *A. tumefaciens* GV3101 carrying pEarleyGate 101::*eveD* and a construct encoding the endosomal marker DsRed-FYVE. Confocal microscopy revealed no colocalisation between EveD-YFP and DsRed-FYVE, suggesting that EveD does not accumulate in endosomal compartments (Fig. 7C).

**Fig. 7.**
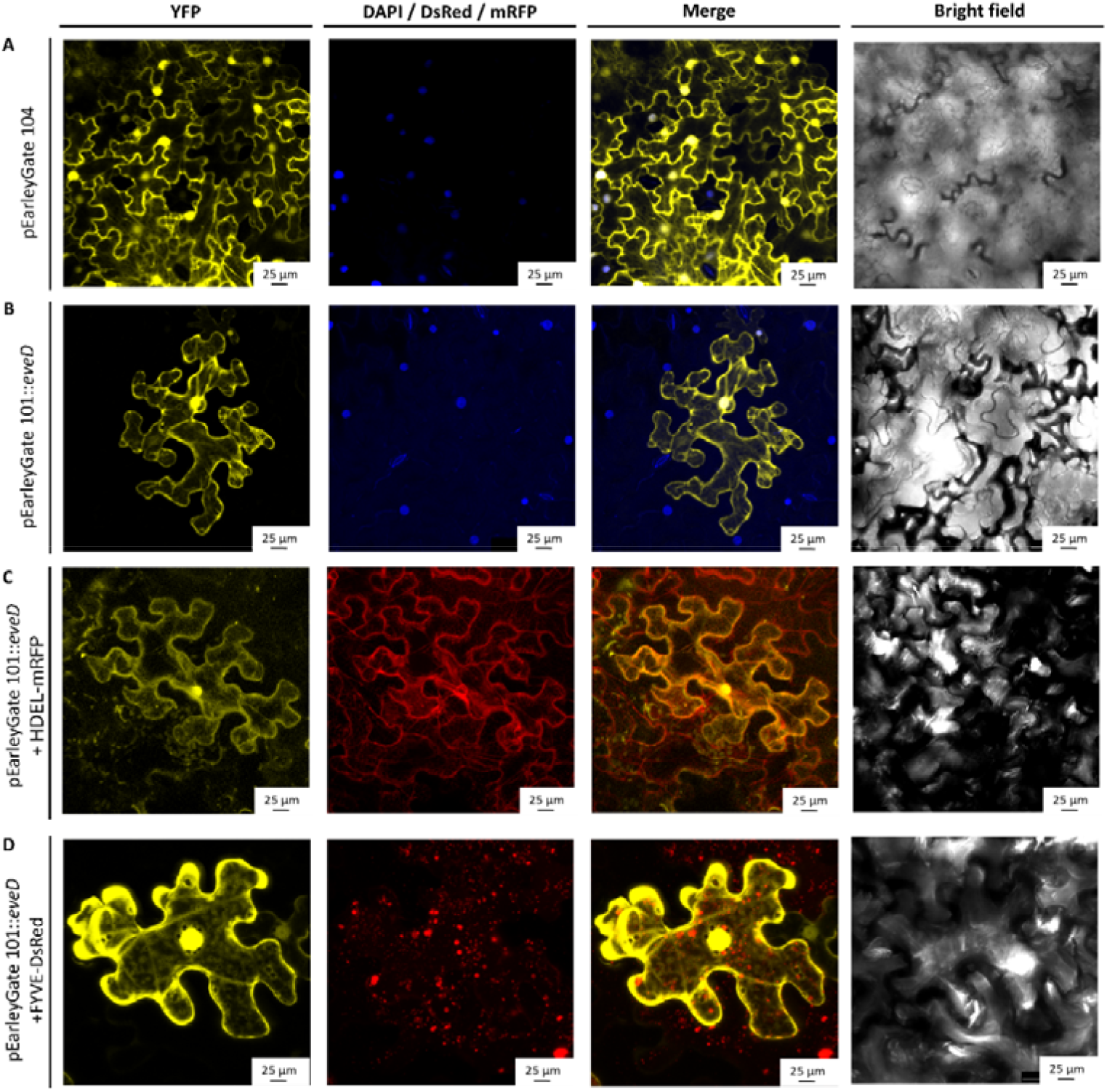
Subcellular localization of *S. fredii* HH103 EveD in *N. benthamiana* leaf cells. Confocal images of *N. benthamiana* leaves 48 h after agroinfiltration. Plasmid pEarleyGate 104 was used as a control for YFP expression (**A**). EveD localisation is shown in yellow (pEarleyGate 101::eveD). Nuclei are shown in blue (DAPI staining) (**B**), whereas the endosome marker FYVE-DsRed (**C**) and the endoplasmic reticulum marker mRFP-HDEL (**D**) are shown in red. Scale bars indicate 25 µm. *Agrobacterium*-mediated transient expression of EveD and subsequent subcellular localisation in *N. benthamiana* leaves were assessed in two independent experiments.

In summary, the results indicate that EveD localises predominantly in the cytoplasm of plant cells, with occasional nuclear accumulation, and associates with the endoplasmic reticulum. This subcellular distribution indicates where EveD accumulates when expressed within plant cells and is compatible with, but does not demonstrate, delivery of EveD into host cells during the interaction.

### EveD Modulates Competitive Fitness and Host-Dependent Symbiotic Interactions of S. fredii HH103

In order to investigate the contribution of EveD to the symbiotic interaction between *S. fredii* HH103 and soybean, nodulation assays were initially performed using the compatible host *G. max* cv. Williams. Plants were inoculated with the parental strain *S. fredii* HH103, the *eveD* mutant, and the complemented strain, and symbiotic parameters were analysed six weeks after inoculation. No significant differences were observed between the parental strain and the *eveD* mutant for any of the analysed parameters, including nodule number, nodule fresh weight, average nodule weight, or shoot dry weight. Similarly, no differences were detected between the complemented strain and the parental strain, confirming that disruption of *eveD* does not affect the overall nodulation capacity or symbiotic efficiency of *S. fredii* HH103 with *G. max* cv. Williams under single-strain inoculation conditions (Supplementary Fig. S9).

Although no evident symbiotic phenotype was observed under individual inoculation conditions, EveD may act at an earlier stage of the symbiotic interaction, with its association with bEVs potentially contributing to this early role. To test this possibility, competition assays were performed between the parental and the *eveD* mutant strains to determine their relative ability to occupy nodules. For this purpose, germinated seeds of *G. max* cv. Williams were inoculated with mixtures of both strains at three different parental strain:mutant ratios (1:1, 10:1 and 1:10), and the proportion of nodules occupied by each strain was subsequently determined. The results revealed a competitive advantage of the *eveD* mutant over the parental strain (Fig. 8A). In the 1:1 inoculation ratio, where an equivalent nodule occupancy would be expected if both strains displayed similar competitive abilities, the mutant strain occupied 58.67% of the nodules (72.94% when mixed nodules were considered), indicating a deviation from the expected 50:50 distribution. Similarly, in the 1:10 inoculation ratio, the mutant occupied 94.27% of nodules (97.21% when mixed nodules were considered), exceeding the expected occupancy of 90%. Conversely, the 10:1 inoculation ratio resulted in a mutant occupancy close to the expected value, with the mutant strain occupying 7.49% of nodules (19.74% when mixed nodules were considered). These results indicate that EveD influences the competitive performance of *S. fredii* HH103 during the early stages of the symbiotic interaction, although it is not required for effective nodulation when the strain is inoculated individually.

**Fig. 8.**
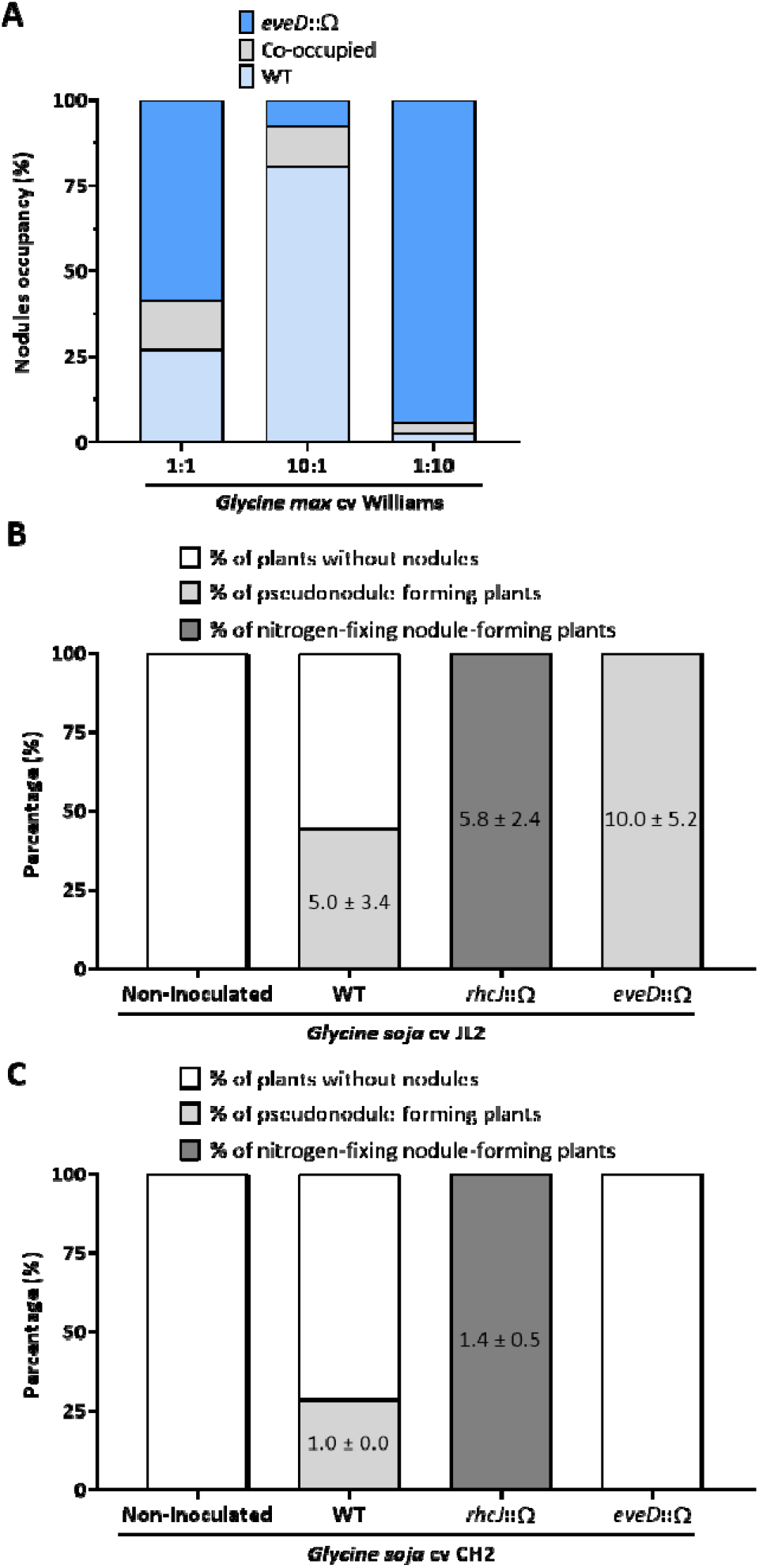
Soybean responses to inoculation with the *S. fredii* HH103 *eveD* mutant. (**A**) *G. max* cv Williams responses to co-inoculation with different *S. fredii* HH103 strains. Percentages of nodule occupancy were evaluated by co-inoculation at different ratios of wild-type (light blue) and *eveD* (blue) strains at 30 days post-inoculation (grey, nodules occupied by both strains). *G. soja* cv JL2 (**B**) or CH2 (**C**) responses to inoculation with the wild-type, *rhcJ* and *eveD* mutant strains of *S. fredii* HH103. Percentages of plants developing nitrogen-fixing nodules (dark grey) or pseudonodules (light grey) were evaluated at 30 days post-inoculation. The mean and standard deviation of the number of nodules and pseudonodules per plant are indicated. Each assay included ten plants per treatment and was performed twice with equivalent results. One representative assay is shown.

Given the contribution of bacterial protein secretion system-associated factors, particularly the T3SS, to host specificity and compatibility in rhizobia, we next investigated whether EveD could influence the interaction of *S. fredii* HH103 with soybean genotypes that are incompatible with this strain. For this purpose, nodulation assays were performed using two wild soybean (*G. soja*) accessions, JL2 and CH2, which are not naturally nodulated by the parental strain *S. fredii* HH103. Notably, previous studies have shown that the incompatibility of *S. fredii* HH103 with these two *G. soja* accessions can be overcome by mutations affecting the T3SS structure or the T3SS regulator TtsI, suggesting that T3SS-associated T3E contribute to host restriction in this specific interaction (Temprano-Vera *et al*., 2018). Our results indicate that, in *G. soja* accession JL2, plants inoculated with the *eveD* mutant strain developed a significantly higher number of pseudonodules compared with plants inoculated with the parental strain *S. fredii* HH103 (Fig. 8B). In contrast, in *G. soja* accession CH2, pseudonodules were detected in plants inoculated with the parental strain, whereas no pseudonodules were observed in plants inoculated with the *eveD* mutant strain (Fig. 8C).

The observed phenotypes suggest that EveD modulates the interaction between *S. fredii* HH103 and soybean in a host genotype-dependent manner, influencing both competitive performance in a compatible host and the outcome of interactions with non-compatible *G. soja* accessions.

## Discussion

The establishment of a successful symbiotic interaction between legumes and rhizobia relies on a complex molecular dialogue involving the reciprocal recognition of both partners and the continuous exchange, perception and integration of a diverse array of signals. Since the pioneering studies that first described the association between rhizobia and legumes, considerable progress has been made in our understanding of the molecular mechanisms underlying this symbiotic interaction (Oldroyd *et al*., 2011; Roy *et al*., 2020). For many years, the main actors considered to mediate this interaction have been flavonoids released by legume roots and NFs synthesised and secreted by rhizobia in response to these compounds, together with other bacterial components such as surface polysaccharides and proteins delivered through different secretion systems, particularly the T3SS (López-Baena *et al*., 2016). However, an important piece of the puzzle may have been overlooked.

Initially regarded mainly as a mechanism for cellular waste disposal, bEVs are now recognised as mediators of intercellular communication and cargo delivery (Kulp and Kuehn, 2010; Toyofuku *et al*., 2019). In plant-microbe interactions, bEVs have been studied predominantly in pathogenic contexts, particularly in relation to virulence and modulation of plant immunity (Janda and Robatzek, 2022; Zhang *et al*., 2024), whereas their contribution to beneficial interactions, including symbiosis, has only recently begun to draw attention. Thus, it has been recently described that rhizobial bEVs can transport NFs and other molecules involved in host recognition and are abundant at the symbiotic interface within root nodules (Taboada *et al*., 2019a; Li *et al*., 2022; Ayala-García *et al*., 2025). These findings point out bEVs may represent an additional and previously underappreciated layer of rhizobium-legume communication, with the potential to transport bacterial molecules beyond those traditionally associated with the establishment and maintenance of symbiosis.

Proteins are of particular interest among the molecular components of rhizobial bEVs because they may include potential effectors involved in bacterial-host communication. Proteomic analyses of *Rhizobium etli* CE3 bEVs identified diverse proteins associated with processes such as adhesion, polysaccharide modification, transport and cell-surface remodelling (Taboada *et al*., 2019a), while a separate study showed that the bEV proteome changes markedly in response to the *nod* gene inducer naringenin (Taboada *et al*., 2019b). Similarly, genistein induction altered the protein and lipid composition of *S. fredii* HH103 bEVs, and more recent analyses showed that bEV protein cargo at the symbiosome varies according to both rhizobial strain and host plant (Li *et al*., 2022; Ayala-García *et al*., 2025). Thus, these findings highlight bEV-associated proteins as a dynamic component of rhizobial-host communication.

Proteins associated with bacterial secretion systems have also been identified in bEVs from different bacterial species (Solé *et al*., 2015; Rybak and Robatzek, 2019; McMillan and Kuehn, 2023; Sirisaengtaksin *et al*., 2023). Notably, several proteins normally associated with T3SS-mediated secretion have been shown to associate with bEVs independently of a functional T3SS, and bEVs can deliver these proteins to host cells, such as SipA, SipC and SopE2 from *Salmonella enterica* serovar Typhimurium, or the T3E translocated intimin receptor (Tir) from enterohaemorrhagic *Escherichia coli* O157 (Kim *et al*., 2019; Sirisaengtaksin *et al*., 2023). These observations support the concept that bEVs can provide an alternative route for the extracellular delivery of effector proteins. In rhizobia, it remains to be determined whether the specific protein cargo of bEVs might also include effectors secreted via specialised secretion systems. In this context, EveD may represent a related, but distinct, case.

Although EveD was initially investigated as a putative T3E based on its similarity to AvrBs3, our experimental analyses revealed a different secretion pattern. EveD is detected in the bEV-enriched fraction, but not in the corresponding bEV-depleted supernatant or in the bacterial cellular fraction under the conditions tested, and its secretion is independent of the T3SS. This distribution argues against its release as a soluble extracellular protein. Proteins exposed at the vesicle surface, such as HtrA and VacA, have been recovered both from vesicles and from the corresponding bEV-depleted medium (Ricci *et al*., 2005; Bartolini *et al*., 2013), whereas EveD is confined to the vesicle fraction. This difference argues against a loosely surface-associated form, but does not establish whether EveD is enclosed within the bEV lumen or anchored at the vesicle surface. Both topologies remain compatible with its distribution across fractions.

Two further observations tie the extracellular pool of EveD to bEV production rather than to its own synthesis. First, *eveD* transcription is essentially invariant across media, growth phases, genetic backgrounds and plant signals, whereas the amount of extracellular EveD differs markedly between non-inducing and inducing conditions. Second, the genistein treatment that increases extracellular EveD also increases vesicle output in this strain (Moreno de Castro *et al*., 2026), hence the extracellular pool of EveD parallels vesicle abundance, not transcript abundance. Given the absence of predicted Sec- or Tat-dependent targeting signals, and since none of the other secretion systems encoded by *S. fredii* HH103 provides a plausible account of its export, the mechanism by which this large soluble protein becomes associated with bEVs remains unknown. One possibility is suggested by the dual origin of bEVs in Gram-negative bacteria: besides blebbing of the outer membrane, which packages periplasmic cargo, vesicles can also arise through explosive cell lysis, generating outer-inner membrane vesicles that entrap cytoplasmic content (Turnbull *et al*., 2016; Toyofuku *et al*., 2019; 2023). Such a route would not require EveD to be translocated across the inner membrane by a dedicated transporter. It would, however, require an intracellular pool of EveD that is not detected in the cellular fraction under any of the conditions tested, which would imply a low intracellular steady-state level. The vesicle population that carries EveD cannot be assigned with the present data.

The regulation of *eveD* does not follow the canonical flavonoid-NodD1-TtsI pathway controlling T3SS gene expression in *S. fredii* HH103. This observation is consistent with genome-wide transcriptomic profiling of the *S. fredii* HH103 genistein stimulon, which revealed a broader flavonoid-responsive gene set that includes genes lacking *nod* or *tts* boxes and regulated independently of NodD1 and TtsI, among which *eveD* was not included (Pérez-Montaño *et al*., 2016). In general terms, our β-galactosidase activity assays and qRT-PCR analyses were consistent with previously published transcriptomic data (Supplementary Table S3; Pérez-Montaño *et al*., 2016; Acosta-Jurado *et al*., 2020; Navarro-Gómez *et al*., 2023). Intriguingly, sequence analysis identified a functional SyrM box in the *eveD* promoter, and *syrM* inactivation resulted in a partial reduction of *eveD* expression, supporting a positive regulatory role for SyrM in *eveD* transcription. Although *syrM* transcription is classically known to be activated by NodD1 upon genistein perception (Acosta-Jurado *et al*., 2020), our findings suggest that *eveD* regulation by SyrM may also occur through a NodD1-independent pathway. Moreover, the reciprocal negative cross-regulation between SyrM and TtsI, whereby each regulator represses the expression of the other (Navarro-Gómez *et al*., 2023), adds a further layer of regulatory complexity to the control of *eveD* and related symbiotic processes in *S. fredii* HH103.

Diverse works have established that TtsI exerts functions outside T3SS activation, modulating surface motility, flagellar gene expression and other cell envelope-associated processes (Alías-Villegas *et al*., 2022; Navarro-Gómez *et al*., 2024; Jiménez-Guerrero *et al*., 2025). While *eveD* transcript levels remained unchanged in a *ttsI* mutant background, EveD is strongly secreted in bEVs in this mutant even in the absence of genistein, a phenomenon not observed in the parental strain or in the corresponding *rhcJ* mutant. These results suggest that, although TtsI levels are low under non-induced conditions, basal TtsI activity might contribute to cell envelope homeostasis independently of T3SS assembly. A precedent exists for a link between the T3SS and vesiculation: in *Pseudomonas syringae* pv. tomato, the *hrcC*, *hrpA* and *hrpL* mutants all produce significantly more bEVs than the wild type when production is normalised to culture density and in the absence of growth differences. This effect was attributed to a reduced number of connections between the inner and outer membranes (McMillan *et al*., 2021), consistent with the general observation that vesiculation rate is set by the density of the links tethering the outer membrane to the peptidoglycan and by the load of material accumulating in the periplasm (Bernadac *et al*., 1998; McBroom and Kuehn, 2007; Schwechheimer *et al*., 2013; Schwechheimer and Kuehn, 2015). In *S. fredii* HH103, however, the increase is observed in the *ttsI* mutant but not in the *rhcJ* mutant, and therefore does not simply follow the loss of a functional secretion apparatus. This points instead to the regulator itself, and remains to be tested directly.

EveD is thus non-canonical in both respects: its transcription is independent of the canonical flavonoid-NodD1-TtsI cascade and its export does not require the T3SS, although TtsI is somehow related. The confirmed T3Es NopC and NopP, by contrast, retain canonical genistein-dependent regulation and are recovered both from the bEV-free supernatant and from the bEV fraction, so in the parental strain the vesicle-associated pool coexists with canonical secretion.

The symbiotic phenotypes observed for the *eveD* mutant indicate that EveD contributes to the interaction of *S. fredii* HH103 with its legume hosts. The phenotype was manifested as an altered competitive behaviour rather than as a change in symbiotic performance upon individual inoculation. Thus, in co-inoculation assays the *eveD* mutant occupied a larger proportion of nodules than the parental strain, indicating that EveD modulates the extent of nodule colonisation without being required for it. A determinant that tempers nodule occupancy is not necessarily disadvantageous for the bacterium, as the number of successful infections is tightly controlled by the host and excessive colonisation is restrained through autoregulation of nodulation (Oldroyd *et al*., 2011). EveD may therefore act as one of the bacterial factors that adjust the level of infection to the host genotype, a role consistent with the genotype-dependent responses observed in the two *G. soja* accessions. Notably, the magnitude of this effect is likely to be underestimated in the co-inoculation assays, since *eveD* mutant cells sharing the rhizosphere with the parental strain are exposed to EveD-containing bEVs released by the latter. Thus, such cross-complementation would make both strains more similar and would therefore reduce, rather than increase, the difference measured between them.

Together with the localisation of EveD in bEVs during free-living growth and its absence from bEVs isolated from bacteroids within soybean or *Lotus burttii* nodules (Ayala-García *et al*., 2025), these findings suggest that bEV-associated EveD may be particularly relevant before or during the early stages of symbiosis, such as root colonisation, host perception or early infection, and not in the establishment of the mature symbiotic state itself.

The T3SS is a major determinant of host range in rhizobium-legume interactions, with translocated T3Es contributing to host restriction by triggering ETI or interfering with symbiosis in non-compatible hosts (Yang *et al*., 2010; Tsukui *et al*., 2013; Jiménez-Guerrero *et al*., 2015b; Yasuda *et al*., 2016; Sugawara *et al*., 2018; Aphaiso *et al*., 2024). Consistent with this, T3SS disruption in *S. fredii* HH103 resulted in nodule or pseudonodule formation in the wild soybean accessions JL2 and CH2, indicating that T3SS-dependent processes constitute a major compatibility barrier in these hosts. In contrast, deletion of *eveD* neither overcame this barrier nor restored the formation of nitrogen-fixing nodules by the host, indicating that EveD is not a primary determinant of T3SS-mediated host restriction. Nevertheless, the distinct phenotypes observed in JL2 and CH2 reveal a host genotype-dependent contribution of EveD to early interaction stages. In JL2, *eveD* deletion increased pseudonodule formation, suggesting that EveD constrains early nodule organogenesis in this background, whereas in CH2 the absence of a functional EveD protein abolished the formation of pseudonodules, indicating that EveD can instead facilitate this response.

Insertion of bEVs into the plant plasma membrane is experimentally established (Tran *et al*., 2022; Rudnicka *et al*., 2022). Endocytic internalisation of intact bacterial vesicles by walled plant cells has not been demonstrated. Because membrane insertion does not require vesicle uptake, both a luminal and a surface-exposed EveD would be brought to the host plasma membrane after bEV-plant cell contact: luminal cargo would be delivered into the cytoplasm, whereas surface-exposed EveD would be deposited at the membrane interface. Either topology is therefore compatible with a function at or beneath the plant plasma membrane.

EveD is a remarkably large protein of 1687 amino acids containing a central region composed of highly conserved tandem repeats, an architecture widespread among proteins that mediate macromolecular interactions (Kajava, 2012). *In silico* structural analyses predict this region to form an extended, predominantly α-helical repetitive architecture, in which tandem repeat units contribute to a continuous α-solenoid-like structure. Although this organisation is reminiscent of the repeated arrays characteristic of TAL effectors, neither sequence- nor structure-based analyses support the classification of EveD as a canonical TAL effector. α-Solenoids constitute a diverse class of repeat-rich protein architectures that can provide interaction surfaces for proteins or nucleic acids and participate in signalling and regulatory processes (Kobe and Kajava, 2000; Fournier *et al*., 2013; Arrías *et al*., 2024), so the extended repeat architecture of EveD may provide a structural basis for interactions with host components, although its specific molecular function remains unknown.

Consistent with this possibility, in the natural legume host EveD is predominantly detected in the cytoplasm, where it appeared associated with structures compatible with the ER, with no apparent association with endosomal compartments, in agreement with its predicted cytoplasmic localisation in eukaryotic cells. EveD is also detected in the nucleus, despite the absence of a canonical NLS. However, the lack of an identifiable NLS does not necessarily preclude nuclear accumulation, as nuclear targeting of microbial effectors can involve mechanisms beyond canonical NLS-dependent import, including interactions with host factors (Harris *et al*., 2023). Thus, the predominant cytoplasmic and ER-associated localisation of EveD, together with its occasional nuclear accumulation, suggests that its repeat-rich architecture could mediate interactions with cytoplasmic or membrane-associated host components and, potentially, with nuclear factors.

Collectively, our findings support a model in which the bEV-associated effector candidate EveD contributes to the early interaction between *S. fredii* HH103 and its legume hosts, highlighting bEVs as an additional route for protein export and a previously underappreciated component of rhizobium-legume communication.

## Acknowledgments

We would like to thank the Spanish Ministry of Science, Innovation and Universities for funding this work through projects PID2022-141156OB-I00, PID2021-122395OA-I00, and PID2025-172427NA-I00. Diego García Rodríguez was supported by a predoctoral contract funded by the University of Seville (VII-PPI US). We thank C. J. Ramos-Rodríguez for performing the genome analysis of *S. fredii* HH103 using PIFAR, and Catherine Jacott for her assistance with *in silico* analyses. We also thank the Microscopy Research Service of the Centro de Investigación, Tecnología e Innovación (CITIUS) of the University of Seville. Some figures were created with BioRender.com. The authors also acknowledge the use of Claude Opus 5 (Anthropic), which assisted in reviewing the manuscript and in improving the clarity of the writing.

## Author Contributions

FJ-LB, and I-JG: conceptualization; JM-V, FJ-LB, and I-JG: methodology; F-PM, FJ-LB, and I-JG: formal analysis; D-GR, P-AG, S-AJ, and F-O: investigation; JM-V, F-PM, JM-B, FJ-LB, and I-JG: resources; F-PM, FJ-LB, and I-JG: data curation; F-PM; FJ-LB, and I-JG: writing - original draft; F-PM, F-O, JM-V, JM-B, FJ-LB, and I-JG: writing - review & editing; F-PM: visualization; FJ-LB, and I-JG: supervision; FJ-LB, JM-V, JM-B, and I-JG: funding acquisition

## Data Availability

All data supporting the findings of this study are included in the manuscript and its supplementary data. The EveD sequence is available in GenBank/ENA under accession CCE94774.1 (locus tag *SFHH103_00272*; *S. fredii* HH103 assembly GCA_000283895.2). Bacterial strains and plasmids generated in this study are available from the corresponding authors on request.

