## Supplementary material for "EveD, a vesicle-associated rhizobial protein shaping nodulation competitiveness and host genotype responses": Supp Information

**This file includes:**

Figures S1 to S9

Tables S1 to S4

Supplementary Dataset S1

Supplementary References

### **Supplementary Figures**

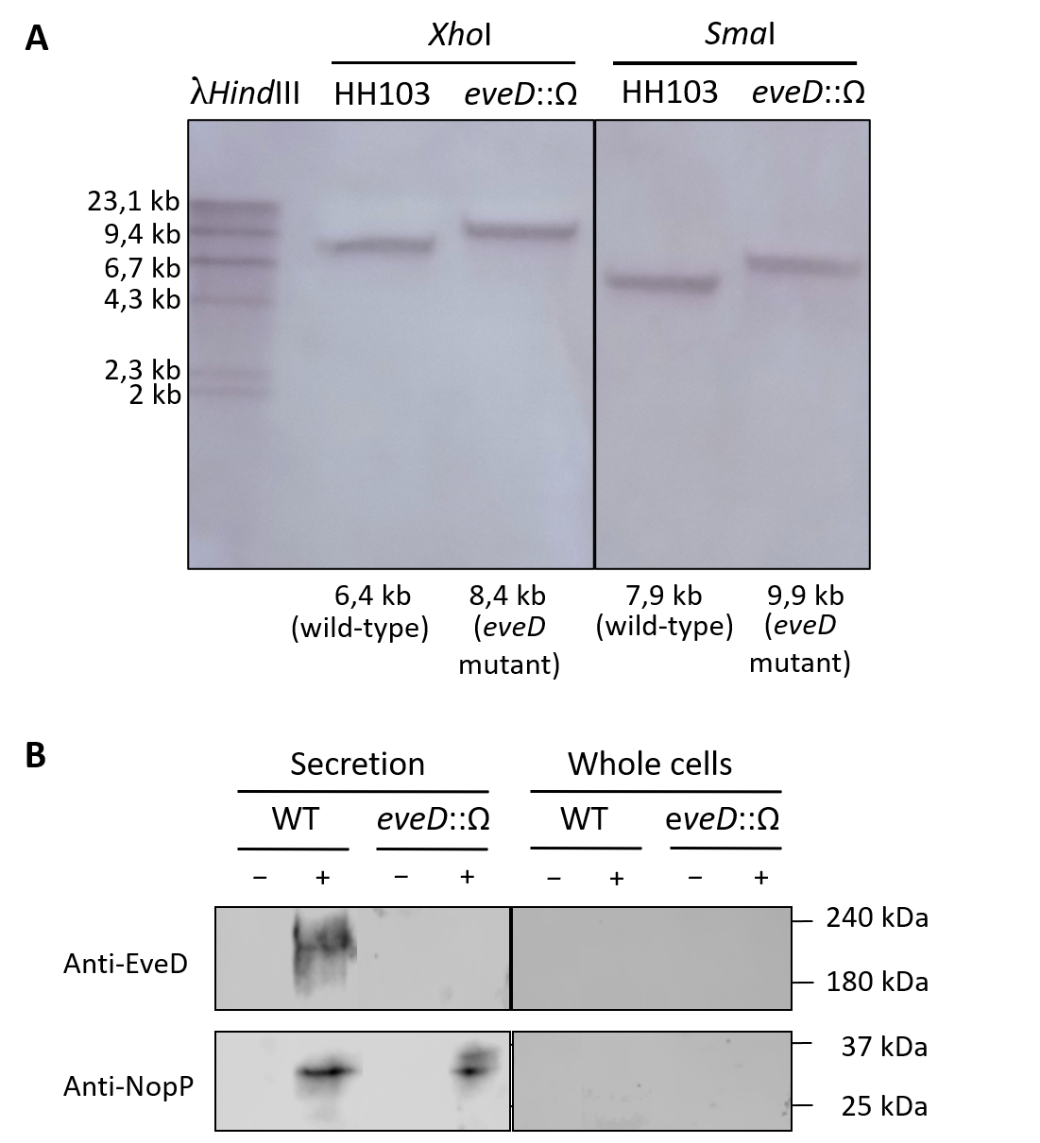

**Fig. S1.** Verification of the mutation of the *S. fredii* HH103 *eveD* gene. (**A**) Southern blot hybridization using an *eveD* gene probe (amplified using primers listed in Table S2). The gDNA of each HH103 strain was digested with *Xho*I or *Sma*I restriction enzymes. (**B**) Production and secretion of EveD and NopP in the parental and *eveD* mutant strains grown in MM medium supplemented (+) or not (-) with genistein. Proteins were detected by Western blot analysis using specific EveD and NopP antibodies. “Whole cells” expression represents the intracellular protein fraction, whereas “Secretion” displays the extracellular protein fraction. The position of the molecular size marker (in kDa) is indicated.

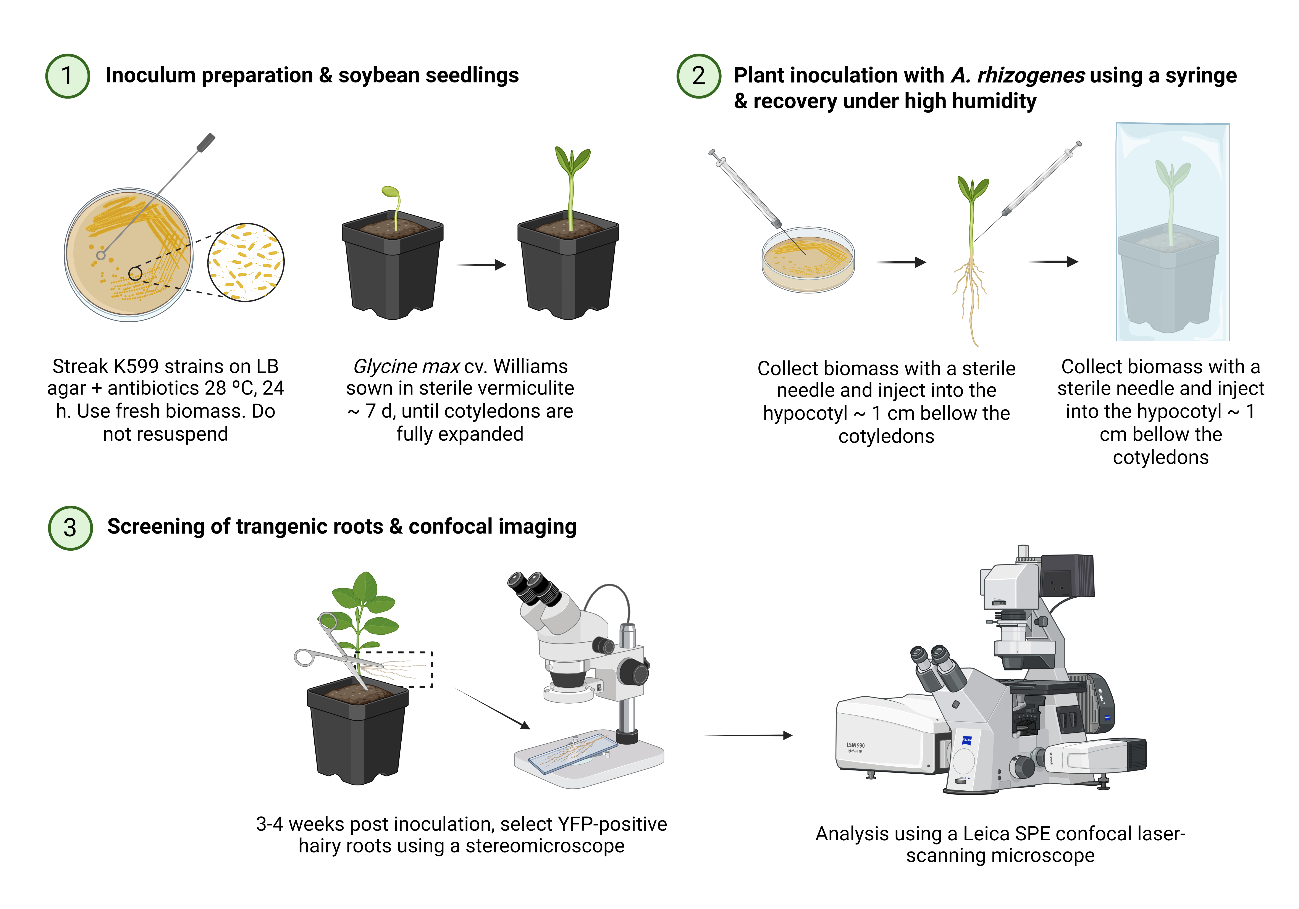

**Fig. S2.** *Agrobacterium rhizogenes* K599-mediated hairy-root transformation of soybean used for the subcellular localisation of EveD. (**1**) Preparation of the K599 inoculum on LB agar with the appropriate antibiotics (28 °C, 24 h) and growth of *Glycine max* cv. Williams seedlings in sterile vermiculite for about 7 days, until the cotyledons are fully expanded; (**2**) inoculation by injection of bacterial biomass into the hypocotyl approximately 1 cm below the cotyledons, followed by recovery under high humidity; (**3**) selection of YFP-positive hairy roots three to four weeks after inoculation using a stereomicroscope, and analysis by confocal laser-scanning microscopy. Created with BioRender.com.

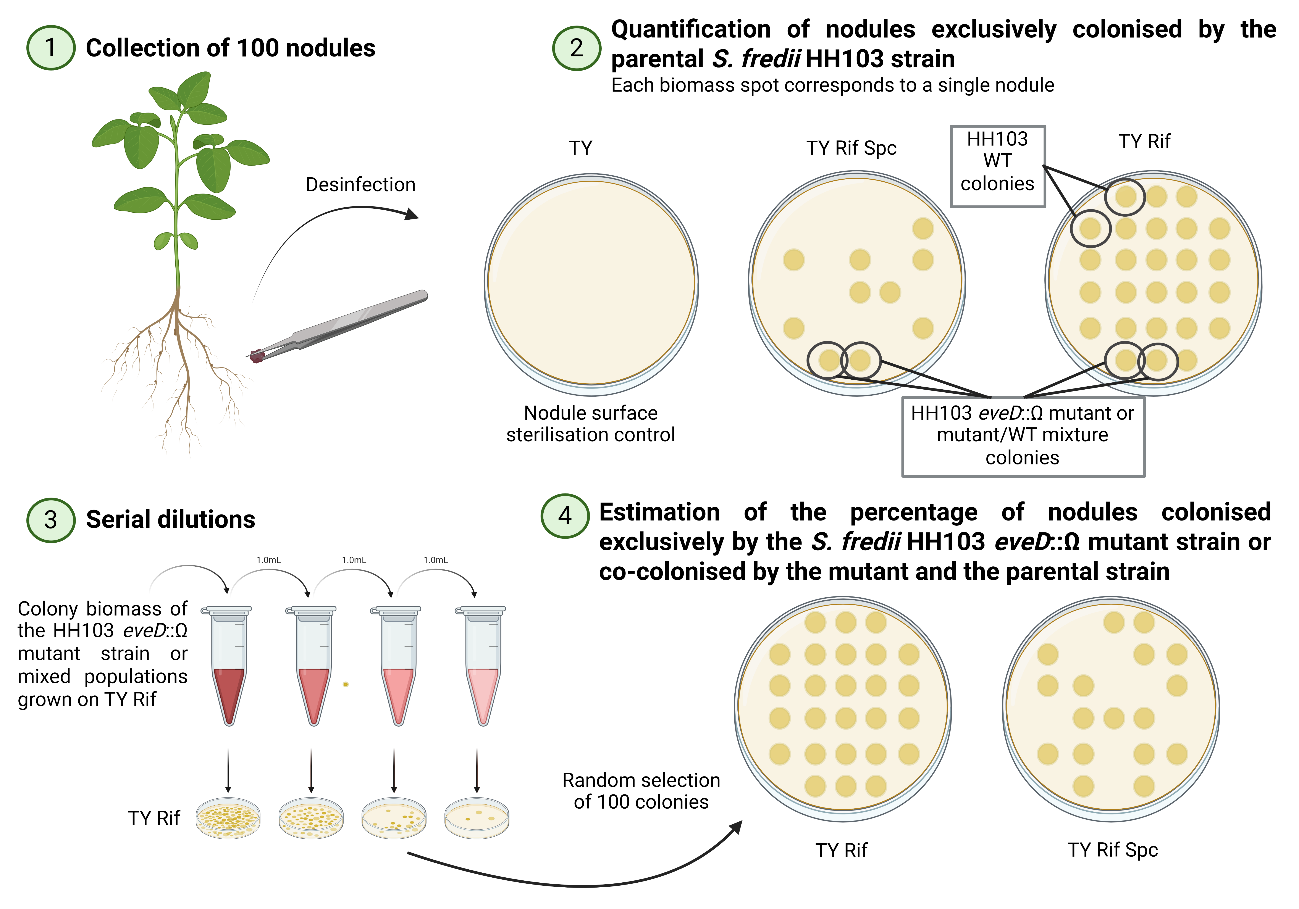

**Fig. S3.** Scheme of the nodule occupancy assay used to assess the competitiveness of the *S. fredii* HH103 *eveD* mutant. (**1**) One hundred nodules per treatment were collected and surface-sterilised; (**2**) nodules occupied exclusively by the parental strain were identified as those yielding growth only on TY+Rif, each biomass spot corresponding to a single nodule; (**3**) biomass recovered from nodules growing on both media was resuspended and serially diluted; (**4**) one hundred randomly selected colonies were replica-plated onto TY+Rif and TY+Rif+Spc to distinguish nodules occupied exclusively by the mutant from nodules co-occupied by both strains. Created with BioRender.com.

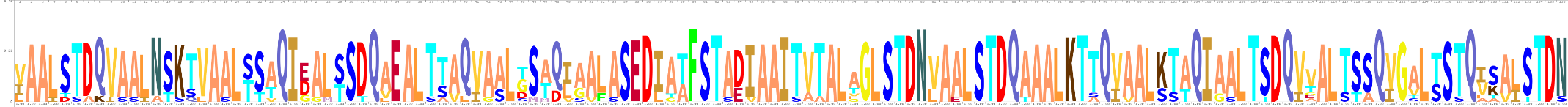

#### **Fig. S4.** Sequence logo of the *S. fredii* HH103 EveD repeat units. The logo was generated with Skylign based on multiple alignments of eleven highly conserved repeat units of 136 amino acids spanning the central region of the protein. Stack height is representative of conservation at that position in the alignment.

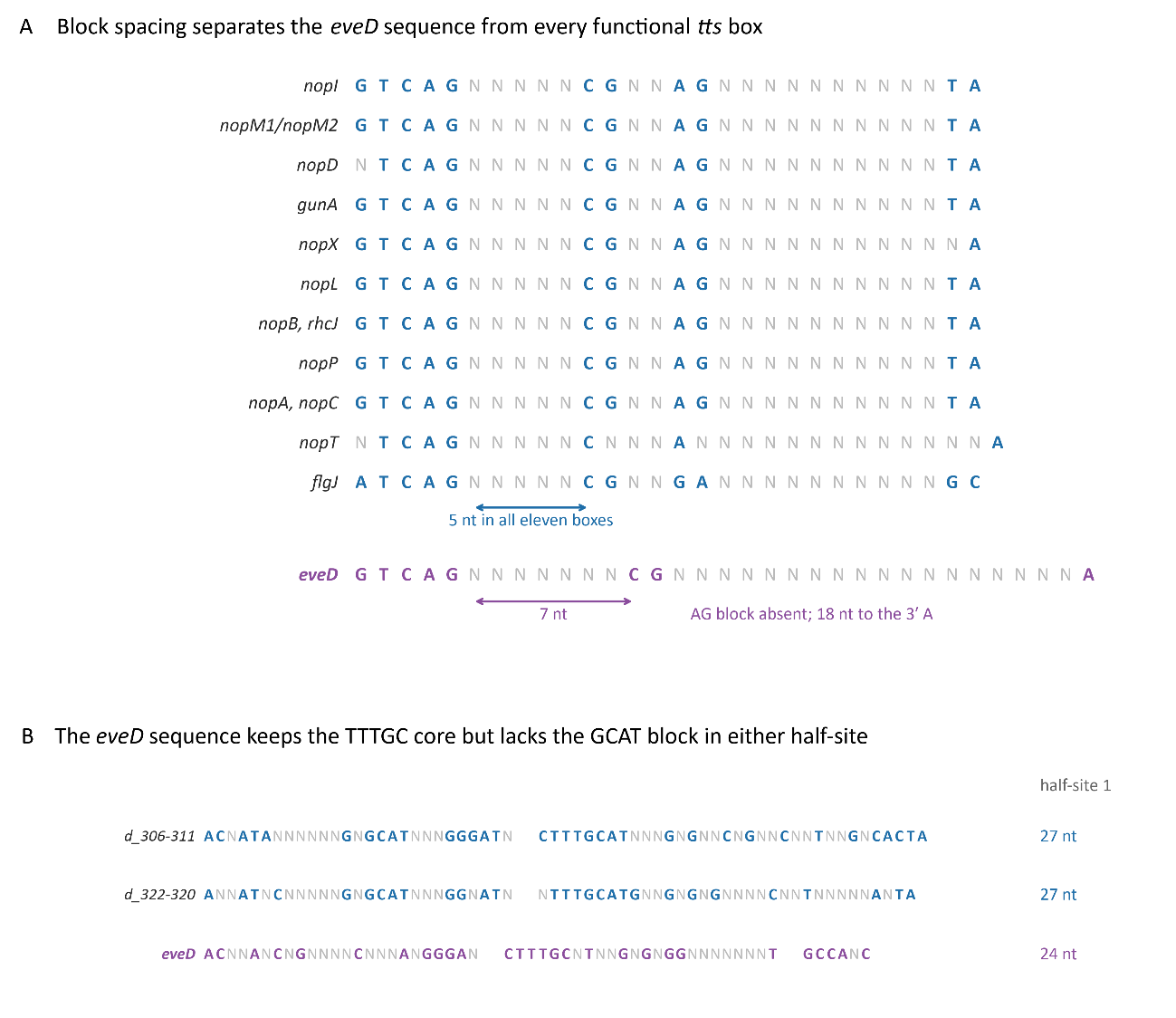

**Fig. S5.** Consensus sequences of the functional *tts* (A) and SyrM (B) regulatory boxes in *S. fredii* HH103 and identification of the corresponding putative boxes in the *eveD* promoter region.

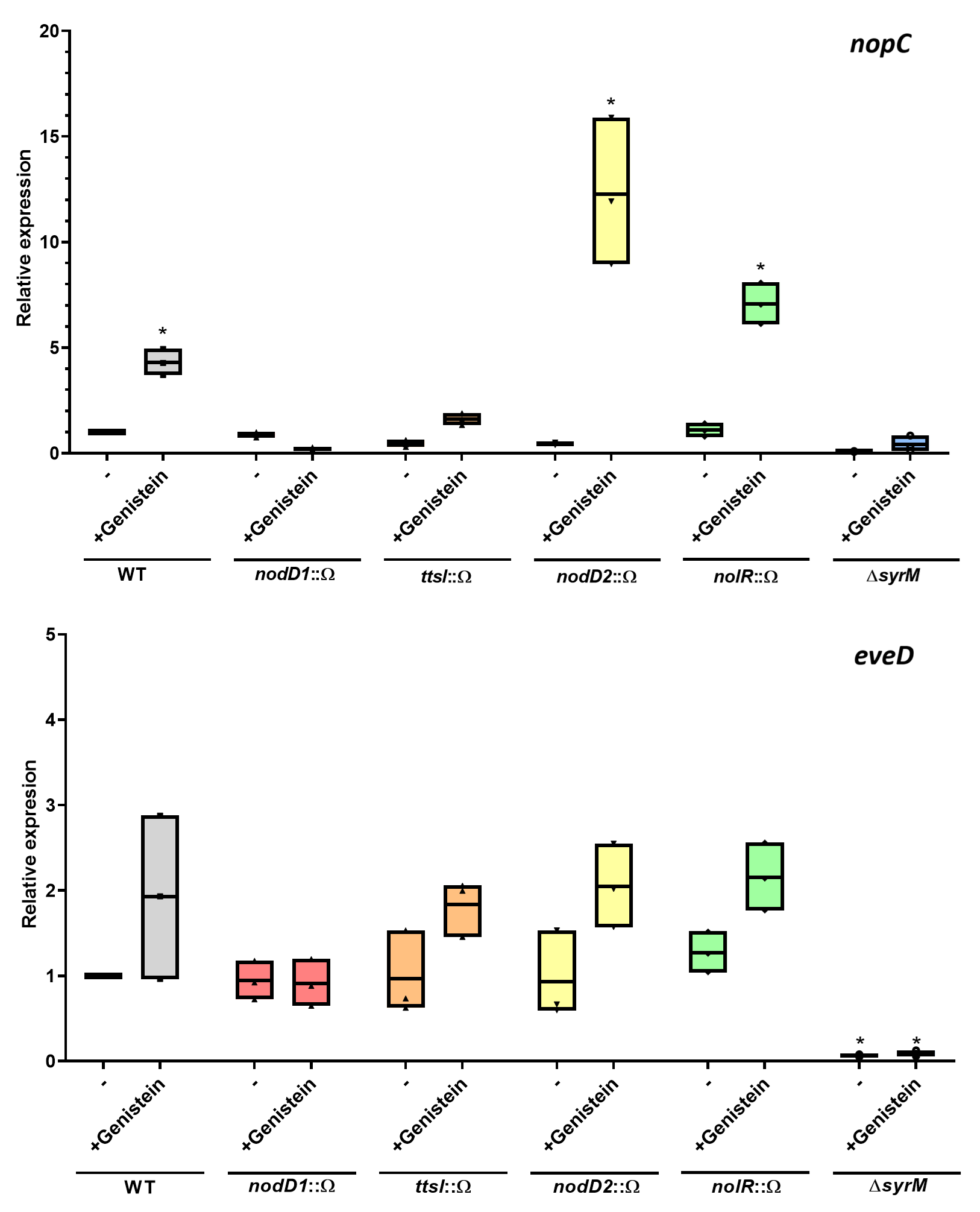

**Fig. S6.** Quantitative real-time PCR analysis of different *S. fredii* HH103 genetic backgrounds. Expression of the different *nopC* (**A**) and *eveD* (**B**) genes was calculated relative to the expression without flavonoids of the parental strain using the Mann-Whitney non-parametrical test. The asterisks indicate a significant difference at the level α = 5% (*P* < 0.05). Parental strain (grey), and *nodD1* (red), *ttsI* (orange), *nodD2* (yellow), *nolR* (green) and *syrM* (blue) strains were grown in MM medium in the presence or absence of genistein. Expression data are presented as the mean (± standard deviation). Three biological replicates were performed, each with six technical replicates. A representative experiment is shown.

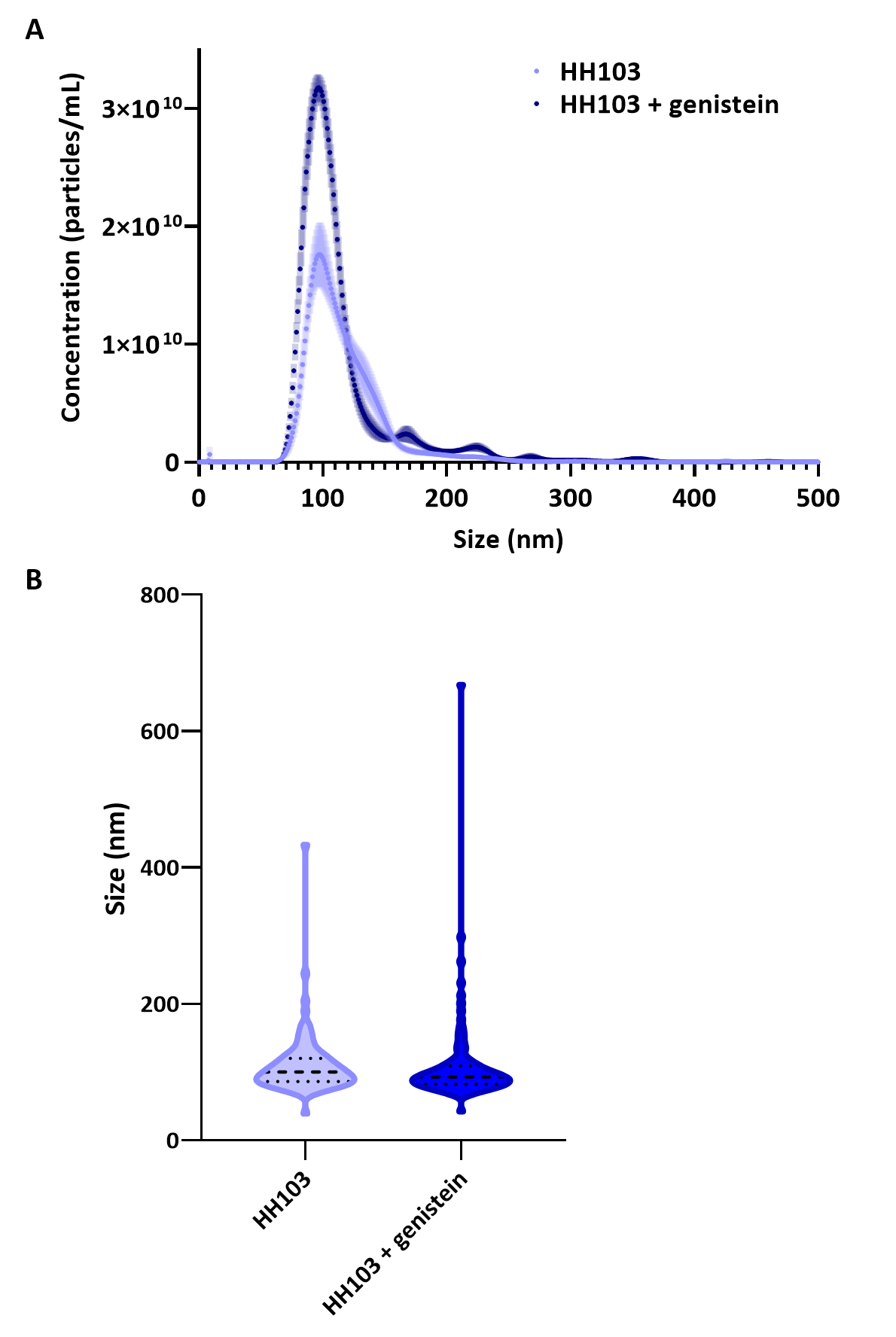

**Fig. S7.** Analysis of bEVs isolated from *S. fredii* HH103. (**A**) Particle concentration (particle/mL) profiles obtained by NTA of bEVs isolated from cultures of *S. fredii* HH103 (HH103) supplemented (dark blue) or not (light blue) with genistein, showing particle concentration as a function of vesicle size. The shaded regions around the curves represent the variation among biological replicates. (**B**) Violin plots summarizing vesicle size distributions obtained by NTA for bEVs derived from cultures of *S. fredii* HH103 (HH103) supplemented (dark blue) or not (light blue) with genistein. Three independent experiments were performed for each condition.

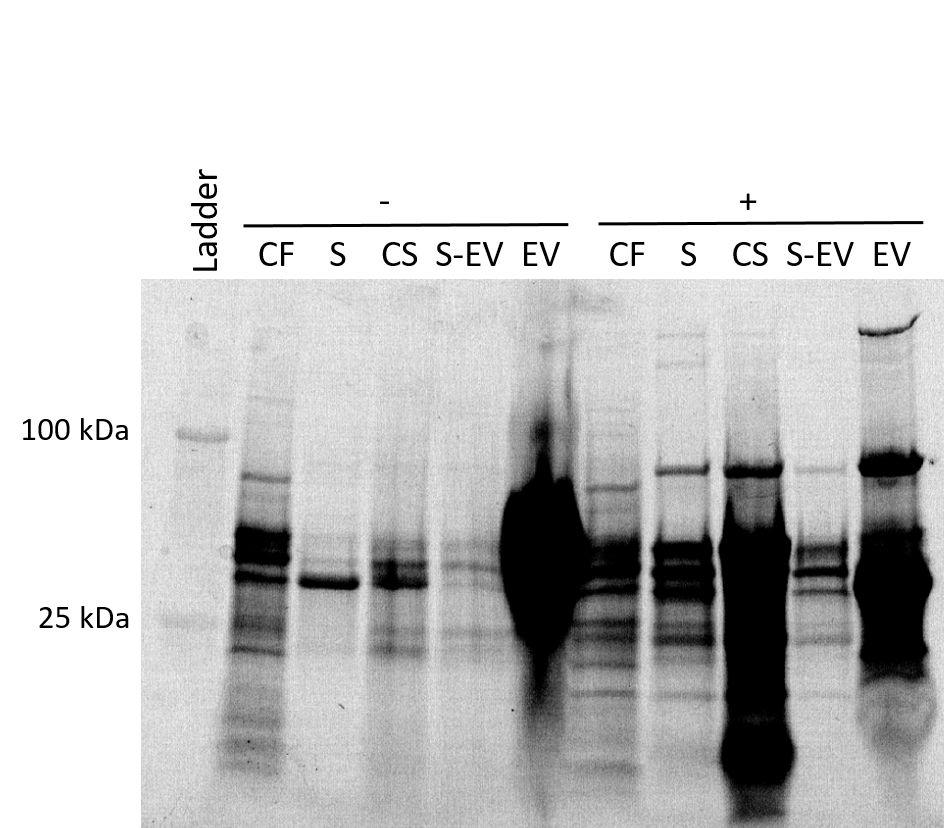

**Fig. S8.** Total protein visualization of different *S. fredii* HH103 fractions. Samples were separated by 15% SDS-PAGE in a stain-free acrylamide gel (Biorad). Following electrophoresis, proteins within the gel were photoactivated and visualized using the Molecular Imager Gel Doc XR+ with Lab Software (Biorad). Bacterial cultures were grown in the absence (-) or presence (+) of genistein. Molecular masses (kDa) of the marker are shown on the left. CF: cellular fraction, S: filtered supernatant, CS: concentrated supernatant, S-EV: EV-free supernatant, and EV: EV-enriched fractions.

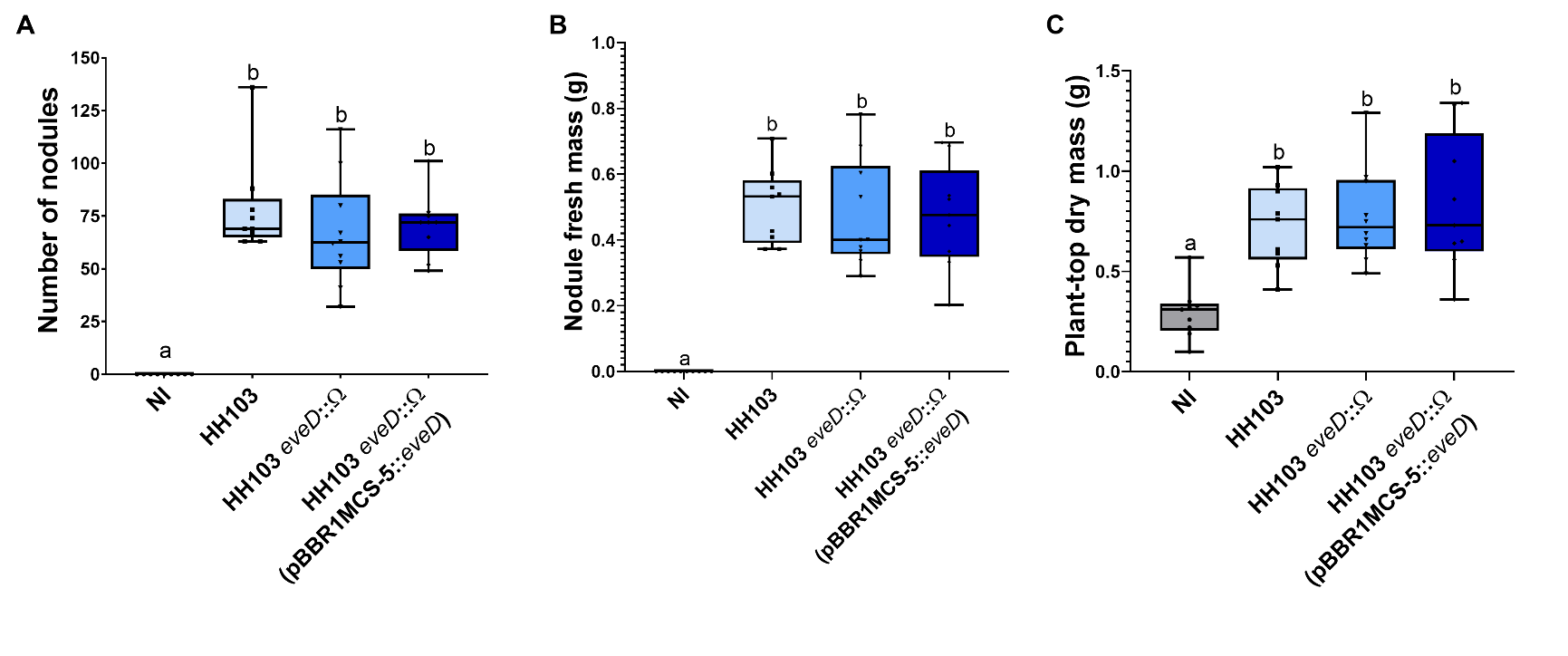

**Fig. S9.** *Glycine max* cv Williams responses to inoculation with *S. fredii* HH103 strains. (**A–C**) Soybean responses to inoculation with different *S. fredii* HH103 strains (WT: light blue, *eveD*::*Ω*: blue, *eveD*::*Ω* (pBBR1MCS-5::*eveD*): dark blue). Plants were evaluated 42 days after inoculation. NI: Non-inoculated plants (grey). Data indicated with the same letter are not significantly different at the level of *α* = 5% (One-Way ANOVA with multiple comparisons, *p* < 0.05). Each assay included ten plants per treatment and was performed twice with equivalent results. One representative assay is shown.

### **Supplementary Tables**

**Table S1.** Bacterial strains used in this study.

| Strain | Relevant properties | Source or reference |
| --- | --- | --- |
| *Sinorhizobium fredii* | | |
| HH103 Rif^R^ | HH103 parental strain; Rif^R^ | Madinabeitia *et al*., 2002 |
| HH103 *nodD1*::Ω | HH103 Rif^R^ derivative containing the Ω interposon inserted into *nodD1*; Rif^R^, Spc^R^ | Vinardell *et al*., 2004 |
| HH103 *nodD2*::Ω | HH103 Rif^R^ derivative containing the Ω interposon inserted into *nodD2*; Rif^R^, Spc^R^ | López-Baena *et al*., 2008 |
| HH103 *nolR*::Ω | HH103 Rif^R^ derivative containing the Ω interposon inserted into *nolR*; Rif^R^, Spc^R^ | López-Baena, 2008 |
| HH103 Δ*syrM* | HH103 Rif^R^ derivative with the *syrM* gene deleted; Rif^R^ | Acosta-Jurado *et al*., 2019 |
| HH103 *ttsI*::Ω | HH103 Rif^R^ derivative containing the Ω interposon inserted into *ttsI*; Rif^R^, Spc^R^ | López-Baena, 2008 |
| HH103 *rhcJ*::Ω | HH103 Rif^R^ derivative containing the Ω interposon inserted into *rhcJ*; Rif^R^, Spc^R^ | Rodrigues *et al*., 2007 |
| HH103 *eveD*::Ω | HH103 Rif^R^ derivative containing the Ω interposon inserted into *eveD*; Rif^R^, Spc^R^ | This work |
| *Escherichia coli* | | |
| DB3.1 | *gyrA462*, *endA1*, ∆(*sr1-recA*), *mcrB*, *mrr*, *hsdS20*, *glnV44* (=*supE44*), *ara14*, *galK2*, *lacY1*, *proA2*, *rpsL20*, *xyl5*, *leuB6*, *mtl1* | Invitrogen |
| DH5α | *supE44*, ∆*lacU169*, *hsdR17*, *recA1*, *endA1*, *gyrA96*, *thi-1*, *relA1*; Nx^R^ | Sambrook *et al.*, 1989 |
| *Agrobacterium tumefaciens* | | |
| GV3101 | Wild-type; Rif^R^ | Rotino and Gleddie, 1990 |
| K599 | Wild-type; Spc^R^ | Savka *et al*., 1990 |
| Plasmid | **Relevant properties** | **Source or reference** |
| pRK2013 | Helper plasmid, ColE1 *ori*; Km^R^ | Figurski and Helinski, 1979 |
| pHP45Ω | Vector containing the Ω interposon, ColE1 *ori*; Spc^R^ | Prentki and Krisch, 1984 |
| pK18*mob* | Cloning vector, suicidal in rhizobia, ColE1 *ori*; Km^R^ | Schäfer *et al*., 1994 |
| pK18*mob*::*eveD* | Plasmid pK18*mob* carrying a ~1.3 kb *Bam*HI-*Xba*I fragment that contains the promoter region and the first 591 bp of HH103 *eveD*, ColE1 *ori*; Km^R^ | This work |
| pK18*mob*::*eveD*::Ω | Plasmid pK18*mob*::*eveD* carrying a 2 kb *Cla*I fragment corresponding to the Ω interposon from pHP45Ω, ColE1 *ori*; Km^R^ | This work |
| pBBR1MCS-5 | Broad host range, multi-copy expression vector, pBBR1 *ori*; Gm^R^ | Kovach *et al*., 1995 |
| pBBR1MCS-5::*eveD* | Plasmid pBBR1MCS-5 carrying a ~5.7 kb *Xba*I-*Bam*HI fragment that contains the promoter region and the full-length *eveD* from HH103, pBBR1 *ori*; Gm^R^ | This work |
| pMP220 | Broad host range, low copy number promoter-probe vector based on the *lacZ* reporter gene (´*lacZ*), RK2 *ori*; Tc^R^ | Spaink *et al*., 1987 |
| pMP220::*P_nopC_* | HH103 *nopC* promoter region cloned into pMP220 as a transcriptional fusion, P*nopC*::*lacZ*, RK2 *ori*; Tc^R^ | This work |
| pMP220::*P_eveD_* | HH103 *eveD* promoter region cloned into pMP220 as a transcriptional fusion, *PeveD*::*lacZ*, RK2 *ori*; Tc^R^ | This work |
| pDONR207 | Gateway donor vector, ColE1 *ori*; Gm^R^ | Invitrogen |
| pDONR207::*eveD* | Plasmid pDONR207 carrying the full-length *eveD* from HH103; ColE1 *ori*, Gm^R^ | This work |
| pEarleyGate 100 | Gateway-compatible plant transformation vector, pVS1 *ori*; Km^R^ | Earley *et al.*, 2006 |
| pEarleyGate 101 | Gateway-compatible plant transformation vector with YFP and HA C-terminal tags, pVS1 *ori*; Km^R^ | Earley *et al.,* 2006 |
| pEarleyGate 101::*eveD* | Plasmid pEarleyGate 101 carrying the full-length *eveD* from HH103, pVS1 *ori*, Km^R^ | This work |
| pEarleyGate 104 | Gateway-compatible plant transformation vector with a YFP N-terminal tag, pVS1 *ori*; Km^R^ | Earley *et al.*, 2006 |
| mRFP-HDEL vector | Plasmid used for transient expression of the endoplasmic reticulum marker HDEL fused to red fluorescent protein, IncP *ori*; Km^R^ | Runions *et al*., 2006; Schoberer *et al*., 2009 |
| DsRed-FYVE vector | Plasmid used for transient expression of the plant endosome marker FYVE fused to the DsRed fluorescent protein, IncP *ori*; Km^R^ | Voigt *et al*., 2005a |

Gm^R^, Km^R^ Nx^R^ Spc^R^, Tc^R^ and Rif^R^ indicate resistance to gentamicin, kanamycin, nalidixic acid, spectinomycin, tetracycline and rifampicin, respectively.

#### **Table S2.** DNA oligonucleotide primers used in this study.

| Name | Sequence^*^ | Use |
| --- | --- | --- |
| eveDext_BamHI | 5'-aatttggatcccgctgagaacatgttcgaatgc-3' | Used to amplify a 5´fragment of HH103 *eveD* including the promoter region for mutagenesis |
| eveDint_omega_XbaIR | 5'-taattatctagagatctacgaagagctcagggc-3' |  |
| InOmega_ClaIF | 5'-ttaattatcgatccggtgattgattgagcaagc-3' | Used to amplify the 2 kb Ω interposon fragment from pHP45Ω for *eveD* mutagenesis |
| InOmega2_ClaIR | 5'-ttattaatcgattgatccggtgattgattgagc-3' |  |
| eveDext_BamHI | 5'-aatttggatcccgctgagaacatgttcgaatgc-3' | Used to amplify the HH103 *eveD* 5´ DNA sequence for *eveD* complementation |
| eveD_IntComp_R | 5'-gagaaggtcgcgatatcttcc-3' |  |
| eveD_IntComp_F | 5'-ccgatcagctcgaagcgctgac-3' | Used to amplify the HH103 *eveD* 3´ DNA sequence for *eveD* complementation |
| eveD_XbaI_HA_R | 5'-atatctagatta**tgcgtaatcaggaacatcgtaagggtatgcgcc**gatggcctggtaggccg-3' |  |
| PeveD_XbaIF | 5'-atatctagaatcccacagacaggcgagttg-3' | Used to amplify the HH103 *eveD* promoter region for *eveD* expression analysis by β-galactosidase |
| PeveD_PstIR | 5'-aaactgcagcatggacgcgagccttcaaag-3' |  |
| PnopC_XbaIF | 5'-atatctagattcgtgtaaagcaagcc-3' | Used to amplify the HH103 *nopC* promoter region for *nopC* expression analysis by β-galactosidase |
| PnopC_PstIR | 5'-aaactgcagcagctactcctgcctta-3' |  |
| eveD_attB1c_F | 5'-*ggggacaagtttgtacaaaaaagcaggct*ccatgacctccatcatctcttctctg-3' | Used for HH103 *eveD Agrobacterium*-mediated transient expression assays |
| eveD_attB2c_R | 5'-*agattggggaccactttgtacaagaaagctgggt*gatggcctggtaggccg-3' |  |
| pDONR_F | 5'-cgttaacgctagcatggatctc-3' | Used for the sequence verification of inserts cloned into pDONR207 |
| pDONR_R | 5'-gtaacatcagagattttgagac-3' |  |
| nopCq_F | 5'-caaaggggggcatgga-3' | Used for qRT-PCR assays |
| nopCq_R | 5'-caaccgatcgaagagcta-3' |  |
| eveDq_F | 5'-agaatctctcgacctctgcc-3' | Used for qRT-PCR assays |
| eveDq_R | 5'-gacatcttccgtatcgagcg-3' |  |
| 16Sq_F | 5'-taaaccacatgctccacc-3' | Used for qRT-PCR assays |
| 16Sq_R | 5'-gataccctggtagtccac-3' |  |

The underlined nucleotides correspond to enzyme restriction sites. The nucleotides in italics correspond to the recombination sequences specific to the Gateway system. The nucleotides in bold indicate the sequence for the addition of an HA epitope sequence.

**Table S3.** Fold change values of selected *S. fredii* HH103 genes in different genetic backgrounds.

|  | **Fold-change^a^** | | | | | |
| --- | --- | --- | --- | --- | --- | --- |
| **Locus tag^b^ / gene** | **WT** | ***nodD1*::Ω** | ***ttsI*::Ω** | ***nodD2*::Ω** | ***nolR*::Ω** | **Δ*syrM*** |
| psfHH103d_126 / *nodA* | 17.65 | -1.38 | 14.68 | 233.98 | 102.56 | 77.22 |
| psfHH103d_3504 / *nopC* | 6.29 | -1.81 | 1.63 | 14.80 | 18.11 | 17.99 |
| SFHH103_00272 / *eveD* | -1.29 | 1.00 | 1.33 | 1.56 | -1.89 | 1.35 |

^a^ Significant transcriptional changes (3-fold induction respect to HH103 noninduced cultures) in the presence of genistein (3.7 µM) is highlighted in light gray. Strong transcriptional changes (30-fold induction) are highlighted in dark gray. RNA-seq data displayed in this table were published by Pérez-Montaño *et al.* (2016); Acosta-Jurado *et al.* (2020) and Navarro-Gómez *et al.* (2023).

**^b^** According to Vinardell *et al.* (2015).

**Table S4.** Candidate secretion routes for EveD in *S. fredii* HH103.

| Route | Evidence in S*. fredii* HH103 | Status |
| --- | --- | --- |
| Sec / Tat | No signal peptide predicted (SignalP-6.0); no transmembrane segment predicted (DeepTMHMM). | Excluded |
| Type I secretion | Two complete systems are encoded (*prtDE*, *SFHH103_03015-03016*, chromosome; *prsDE*, *SFHH103_06228-06229*, plasmid pSfHH103e; Vinardell *et al*., 2015), but both are distant from *eveD* and each is flanked by its own candidate substrate, whereas type I substrates are typically encoded next to their cognate transporter. EveD contains no RTX-type calcium-binding nonapeptide (GGxGxDxUx), whereas the same search recovers 10-29 copies in each of the eight HH103 proteins annotated as haemolysin-type calcium-binding or RTX-like, with which EveD shares no detectable sequence similarity; accordingly, EveD is not among the 18 proteins previously proposed as putative type I secretion substrates in this genome on the basis of a genome-wide GG-repeat search combined with subcellular localisation prediction (Vinardell *et al*., 2015). The only feature compatible with this route is the composition of the C-terminal 60 residues, which are acidic (net charge -8; first percentile of the proteome) and hydrophobic (51.7 %), as in uncleaved type I signals; this is a compositional resemblance rather than a defined motif, and type I secretion cannot be formally excluded on that basis alone. | Unlikely |
| Type II secretion | A complete cluster is encoded on the chromosome (*gspD*-*gspM* together with the prepilin peptidase *gspO*; *SFHH103_02678-02688*; Vinardell *et al*., 2015). Type II substrates must first reach the periplasm through the Sec or Tat pathways, neither of which EveD can use. | Excluded |
| Type III secretion | EveD secretion is unaffected in the *rhcJ* and *ttsI* mutant backgrounds (Fig. 4). | Excluded experimentally |
| Type IV secretion | Three plasmid-borne *virB* clusters are encoded: two complete, on pSfHH103d and on pSfHH103e (*SFHH103_06274-06285*), and one truncated, on pSfHH103c (*SFHH103_04078-04089*) (Vinardell *et al*., 2015). Type IV substrates are translocated by contact-dependent machinery and are not released into the cell-free medium, whereas EveD is recovered from cell-free culture supernatant. | Excluded |
| Type V secretion (Va, Vb, Vc, Ve) | No classical autotransporter, TpsB/FhaC-family transporter, trimeric autotransporter adhesin or intimin/invasin homologue is encoded; the only related gene is the autotransporter assembly factor TamA (SFHH103_03105); no protein similar to known type V transport units was identified in the original analysis of this genome either (Vinardell *et* *al*., 2015). The AlphaFold model of EveD contains no β-strand and therefore no autotransporter β-barrel. | Excluded |
| Type VI secretion | No Tss, Hcp, VgrG, IcmF, ClpV or PAAR component is encoded in the genome, in agreement with the original analysis of the HH103 genome (Vinardell *et al*., 2015) and in contrast to *S. fredii* USDA257 (Reyes-Pérez *et al*., 2025). | Excluded |
| EV-associated release | EveD is recovered in EV-containing fractions and is absent from the EV-depleted supernatant (Fig. 5). | Observed |

Secretion-system components were searched for in the PGAP annotation of assembly GCF_048585425.2 and in the original annotation of assembly GCA_000283895.2, and the resulting inventory was compared with that reported by Vinardell *et al*. (2015). Locus tags follow the SFHH103_ nomenclature of GCA_000283895.2.

**Supplementary Dataset S1.** Sequences used to build the EveD phylogeny (provided as a separate Excel file). The table lists, for each of the 133 tree leaves, the NCBI accession, strain locus tag where available, organism, taxonomic note, protein length, NCBI annotation, clade assignment and its bootstrap support, percentage identity and JTT maximum-likelihood distance to EveD of *S. fredii* HH103, the number of sequences represented by the leaf and the accessions collapsed into it. A second sheet summarises the seven clades.
